# multiVIB: A unified probabilistic contrastive learning framework for atlas-scale integration of single-cell multi-omics data

**DOI:** 10.1101/2025.11.29.691308

**Authors:** Yang Xu, Stephen J. Fleming, Brice Wang, Erin G. Schoenbeck, Mehrtash Babadi, Bing-Xing Huo

## Abstract

Comprehensive brain cell atlases are essential for understanding neural function and advancing translational research. As single-cell technologies proliferate across platforms, species, and modalities, atlas construction increasingly depends on integration frameworks that are capable of aligning heterogeneous datasets without obscuring biological variations. However, existing methods are typically limited to narrow use cases, often requiring *ad hoc* workflows that may introduce artifacts. Here, we introduce multiVIB, a unified probabilistic contrastive learning framework for diverse scenarios of data integration. We demonstrate that multiVIB achieves state-of-the-art performance while minimizing spurious alignments. Applied to atlas-scale datasets generated by the BRAIN Initiative, multiVIB supports robust and scalable integration across data modalities and preserves species-specific variation in cross-species analyses. These results position multiVIB as a scalable, biologically faithful framework for the construction of next-generation brain cell atlases.

## 1 Introduction

The molecular profiles of individual cells underpin the complex neural behaviors, cellular interactions, and regulatory circuits that collectively define the mammalian brain[1]. To decode this logic, the BRAIN Initiative established the Cell Census Network (BICCN) (RRID:SCR 015820) and the succeeding Cell Atlas Network (BICAN) (RRID:SCR 022794) to build comprehensive atlases of the mammalian brains at single-cell level with molecular resolution. While landmark studies have produced high-quality reference atlases for the mouse[2] and primate (Johansen *et al*.[3]) brains using standardized pipelines within a single research center, the broader consortium has generated and shared a far more diverse landscape of single-cell brain datasets, potentially capturing additional biological variations, awaiting incorporation into these central resources[4, 5]. To realize this vision of an inclusive and evolving brain atlas, computational frameworks that can integrate heterogeneous single-cell multi-omics datasets and preserve the biological fidelity are essential. Furthermore, the ultimate goal of constructing these atlases is not only to elucidate cellular and molecular mechanisms of brain, but also to realize their translational potential. This would also require rigorous comparison of the human brain with other model organisms[6], which again necessitates integration tools that are capable of aligning datasets across species as well as preserving the biological specificity that distinguishes species.

A common strategy for integrating heterogeneous single-cell multi-omics datasets is to adapt batch-correction methods, by treating differences of studies, species, or modalities as covariates and attempting to correct them as if they were technical batch effects[7–13]. While effective in some integration tasks, comprehensive benchmarking studies have shown that such approaches frequently collapse distinct cell states and suppresses genuine biological differences[14–18]. Particularly in cross-modal integration, batch-correction-like strategy requires engineering shared features across modalities. However, bridging different modalities through shared features is feasible but not always straightforward[19, 20]. An alternative strategy would rely on joint-profiling technologies[21– 24] that offer a biologically grounded solution where multiple assays were directly probed from the same cells. These jointly profiled multi-omics datasets, therefore, supply paired cells as natural anchors for multimodal integration[25–28]. However, jointly profiled datasets constitute only a small fraction of all single-cell data generated to date and are likely to under-represent the whole cell populations. While jointly profiled cells provide valuable anchors for multimodal integration, limited or unrepresentative anchor sets can produce spurious correspondences across modalities, where unpaired cells in different modalities are incorrectly aligned or modality-specific regulatory programs are collapsed into an artificial consensus[29].

Besides cross-modal integration, cross-species integration is emerging as a key priority in brain atlas construction. Data alignment across species is typically achieved by focusing on conserved homologous genes[15, 30], which may inadvertently eliminate species-specific biology. Some new approaches improved cross-species alignment by drawing on external biological priors such as curated cell-type annotations or pretrained protein language models[31, 32]. However, dependence on these scarce external resources could limit broader application of the tools, as these external resources are either not available in certain species or deployment of these external resources is not scalable.

As large consortia like BICAN expand toward tens of millions of cells across diverse modalities and species, the risk of generating harmonized yet biologically incorrect embeddings jeopardizes the scientific value of next-generation cell atlases. To address these challenges, we developed multiVIB, a biologically grounded and technically versatile framework for integrating single-cell datasets across studies, species, and molecular modalities. Built on the variational information bottleneck (VIB) principle[33] and probabilistic contrastive learning[34, 35], multiVIB learns latent representations that reflect shared biology while preserving species-, modality-, and study–specific variations. We demonstrate the advantages of multiVIB through two atlas-scale case studies: 1) integrating multi-omics datasets of the mouse primary motor cortex from multiple studies into a coherent, scalable atlas framework; and 2) recovering the cross-species consensus taxonomy of the primate basal ganglia while preserving species-resolved regulatory programs. These results show that the probabilistic design of multiVIB effectively accommodates integration over diverse scenarios and successfully preserves biologically meaningful variation amid technical differences.

## 2 Design

Single-cell multi-omics data integration strategies can be organized into two principles[36](Fig. 1A): horizontal integration, which aligns datasets through shared genomic features, and vertical integration, which uses jointly profiled multi-omics cells as direct biological anchors across modalities. Although widely used, neither strategy alone is sufficient for the mosaic structure of real-world scenarios where feature overlap and assay pairing are both partial and inconsistent across studies (Fig. 1A). In these cases, researchers have to select tools and assemble bespoke workflows that can be easily prone to artifacts like spurious alignment. Spurious alignment arises when algorithms over-correct technical differences by collapsing distinct biological features or when disjointed pipelines are applied to complex mosaic datasets for which they were not designed. Note that a third approach, diagonal integration, which attempts to infer correspondences without explicit anchors[36], was excluded from our framework because it is particularly susceptible to spurious matches and is generally unsuitable for atlas-scale tasks[29].

**Figure 1:**
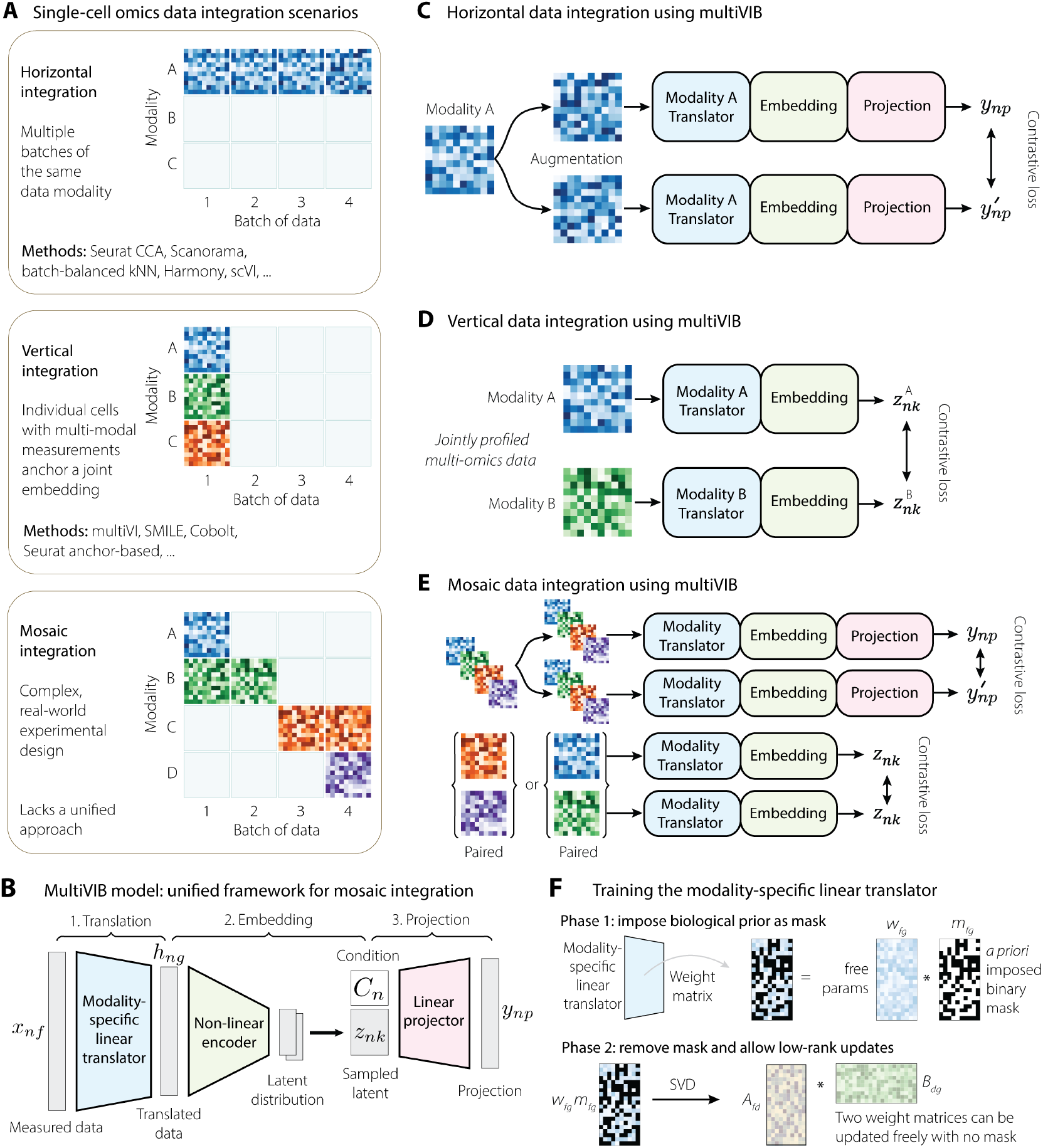
A comprehensive framework for single-cell multi-omics data integration. (A) Three different data integration scenarios are presented. Horizontal integration is the case where there are multiple batches of the same data modality or cross-modal integration through shared features. Vertical integration is the case where individual cells are jointly profiled, collecting several data modalities per cell. Each of these cases have several computational tools available to perform data integration. Mosaic integration combines horizontal and vertical strategies to accommodate more complex data scenarios, particularly when multiple batches of multi-omics datasets are generated across different experiments. In such cases, neither horizontal nor vertical integration alone is sufficient to achieve a comprehensive integration. (B) The components of the multiVIB model. The trainable backbone consists of 3 parts: a modality-specific translator, an encoder, and a projector. Input data *x*_*nf*_ is first translated into a gene-centered data *h*_*ng*_. Encoder then projects the translated data *h*_*ng*_ into latent representation *z*_*nk*_. *z*_*nk*_ is concatenated with batch, species, or modality covariates *C*, and projector embeds the concatenated output into a slightly higher dimension space *y*_*np*_. After training, *z*_*nk*_ becomes the integrated latent representation of cell *n*. (C) Horizontal integration is accomplished by applying data augmentation to a cell in a single modality, paired with a contrastive loss on projected embeddings *y*_*np*_ of two views, where the projector is conditioned on each cell’s batch label *C*_*n*_. (D) Vertical integration is accomplished by applying a contrastive loss on the latent embeddings *z*_*nk*_ of two modalities from the same cell. Training puts positive counterparts 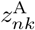 and 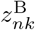 of the same cell close together. (E) Mosaic integration is accomplished using the same framework, with an appropriate data-dependent mix of the horizontal and vertical strategies. (F) The modality-specific linear translator has two training phases in horizontal integration. The first phase imposes a biologically-informed prior as a mask. The second phase relaxes this constraint. Using the phase 1 weight matrix as a starting point, the weight matrix is decomposed into the low-rank matrices *A*_*fd*_ and *B*_*dg*_, which are then updated freely.

multiVIB was designed as a unified probabilistic framework that operates flexibly across horizontal, vertical, and mosaic configurations and produces biologically grounded embeddings. Rather than building separate methods for each integration scenario, multiVIB boasts a single generalizable model backbone, supported by two key principles: (1) uncertainty-aware latent representations, derived through the variational information bottleneck (VIB), which prevent overconfident or forced alignments; and (2) probabilistic contrastive learning (PCL), which reinforces reproducible correspondences across datasets.

At the core of multiVIB is a shared variational encoder that projects all cells, regardless of modality or species, into a common latent space *z*_*nk*_ (Fig. 1B). Each dataset, based on its distinct input feature space, would first go through a modality- or species-specific translator that maps heterogeneous measurements — such as chromatin accessibility, methylation, or gene expression — into a shared space of gene-expression dimensionality. This shared space has the same dimensionality as the gene expression feature space (for example, the number of highly variable genes) and is anchored to gene expression because RNA-seq is the most common molecular readout across multi-omics datasets. The values along each axis after translation are learned representations rather than directly interpretable gene counts. The shared encoder then approximates a stochastic embedding for the translated data to capture biological uncertainty, while a lightweight projector incorporates batch, species, and modality covariates to prevent technical factors from shaping the latent space.

With this backbone fixed, multiVIB adapts to different integration scenarios by altering only the training strategy, not the architecture (Fig. 1C–E). For *horizontal integration* (no jointly profiled cells; Fig. 1C), datasets are anchored through shared features, and multiVIB aligns cells by enforcing consistency across shared genomic signals while ensuring that technical covariates do not drive the alignment. To apply the horizontal approach in cross-modal or cross-species integration, multiVIB requires masking translator with biological-informed prior for reliable data alignment during training (See Methods 5.3.) For *vertical integration* (jointly profiled multimodal cells; Fig. 1D), jointly profiled multi-omics data covers individual cells with multiple modality views. These cells serve as direct biological anchors, allowing multiVIB to learn cross-modal correspondence without relying on engineered feature mappings. *Mosaic integration* (Fig. 1E) is achieved by combining horizontal and vertical steps tailored to the pattern of modality overlap (See Methods 5.1 for further details).

The design of multiVIB, involving VIB and PCL, provides principled safeguards against failure of representation learning that commonly drives spurious alignment. The VIB imposes stochastic and uncertainty-aware embeddings that prevent overconfident alignment, while the contrastive objective reinforces reproducible similarity across datasets. These properties position multiVIB not simply as another option among existing tools, but fundamentally as a more reliable integration backbone. We next show how these design choices lead to stronger and more robust integration outcomes in different integration scenarios.

## 3 Results

### 3.1 multiVIB mitigates spurious alignment in single-cell integration

Spurious alignment, where biologically distinct cell states are incorrectly mapped in the same latent location, is a pervasive challenge in single-cell data integration. Such errors can propagate through downstream analyses and even into the step of constructing reference atlases. To avoid spurious alignment, horizontal integration requires reliable shared genomic features across modalities. In multiVIB, we designed two forms of modality-specific translator to address cross-modal integration under the horizontal paradigm. The first form is a masked linear translation from input-specific feature space to shared-gene intermediate space. This mask is constructed based on a biological prior. Take the translation of chromatin accessibility peak data into a gene expression output as an example, mask informs the model if a chromatin opening region contributes to transcription of a relevant gene. Doing so can prevent the model from exploiting feature correlations solely to fit within-dataset variation. However, adding such mask in training also eliminates recovery of additional cross-dataset covariations. Therefore, we introduce the second form in the way of lower-rank matrix decomposition. Once an initial biologically consistent alignment was achieved through training the translator in the first form, we continue training the model with the second form, which gives the model more freedom to recover additional cross-dataset covariations without breaking the biological coherence (Fig. 1F).

In vertical integration, which relies on limited jointly profiled cells, hidden mismatches or underrepresented cell populations can lead to incorrect cross-modal correspondences that are far more difficult to detect or correct, posing a deeper risk for multimodal integration (Fig. 2A). To illustrate that the design of multiVIB offers a robust solution to alleviate this issue, we set up a conceptual test using 10X Genomics Multiome PBMC data[37], a well-established dataset that includes joint RNA and ATAC measurements at the single-cell level, along with granular cell-type annotations. We deliberately blinded models to the true cell correspondences between paired RNA and ATAC measurements in 3 cell types: CD16+ monocytes, CD8+ T cells, and B1 B cells, but used those correspondences as ground truth for validation. For other cell types, models would see paired cells between the RNA and ATAC measurements to learn correspondence of these two modalities. In this setup, we compared cross-modal alignments by multiVIB and four other vertical integration methods: Cobolt[25], SMILE[26], MultiVI[27], and Seurat (dictionary learning)[28] (RRID:SCR 016341). We observed that multiVIB showed stronger performance in both modality alignment (modality mixing score) and cross-modal mapping (macro F1) (Fig. 2B). Unlike alternative approaches whose performance degraded as latent dimensionality increased, multiVIB remained robust across a broad range of latent dimensions. Using the local annotation enrichment score (LAES)[13], which measures modality mixing in each cell’s neighborhood, multiVIB also achieved the best local-scale alignment across all three blinded cell types (Fig. 2C).

**Figure 2:**
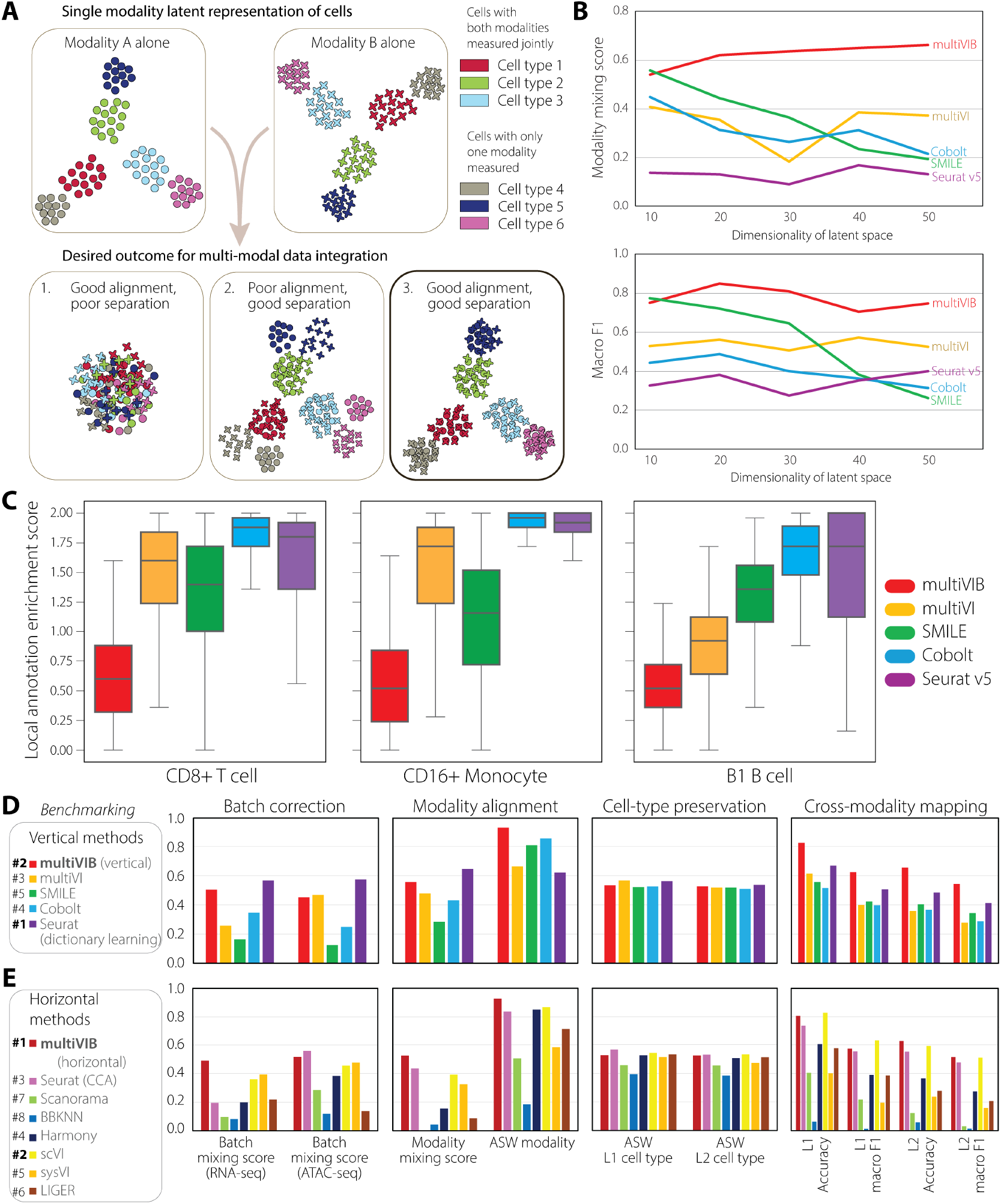
Benchmarking data integration using mutiVIB and existing methods. (A) Conceptual illustration of “spurious alignment” in vertical integration. The thought experiment is that we have a jointly profiled dataset for modalities A and B. In cell types 1–3, jointly profiled data are used during data integration, but for cell types 4–6, the model does not know these are jointly profiled measurements. The joint nature of data collection in cell types 4–6 is withheld and used as ground truth for validation of integration outcome. Data integration can result in various outcomes for cell types 4–6, whose labels are not used during training. Schematic cluster plots show three potential outcomes. “Spurious alignment” would include outcomes 1 and 2, which both are not ideal results. (B) Evaluation of cross-modal alignment in three blinded cell types, similar to the setup in (A), using real 10X Multiome PBMC data. Modality mixing score quantifies the degree of alignment between modalities A and B on the blinded cell types, while label transfer (macro F1) assesses the biological correctness of that alignment. Both metrics are plotted against latent dimensionality to illustrate how embedding dimension also affects integration performance (Higher values in both metrics indicate more successful integration). (C) Local Annotation Enrichment Score (LAES) for the three blinded cell types. LAES quantifies the modality composition of each cell’s nearest neighborhood; lower values indicate better local mixing between two modalities and fewer modality-driven clusters. (D) Benchmarking of 5 vertical integration methods using metrics in 4 categories. (E) Similar benchmarking as (D) but in 6 horizontal methods. L1: coarse cell type annotations; L2: granular cell type annotations. All metrics in all 4 categories are in the range from 0 to 1 and are all annotated as higher is better.

Meanwhile, spurious alignment of horizontal integration also arises from over-correcting technical differences and collapsing distinct cell states into the same latent location. To assess whether multiVIB would incorrectly merge cell populations that are present in only a subset of datasets, we examined its behavior on an expanded dataset that combines PBMC and BMMC scRNA-seq datasets from 4 different studies[14]. In particular, the dataset from Oetjen *et al*. contains progenitor cell types that are not present in the other 3 studies[38]. These study-specific progenitor cells provide a natural test for over-integration. We observed that multiVIB preserved these progenitor populations as distinct groups rather than forcing them to mix with cells from the other studies (Supplementary Fig. S1), demonstrating that multiVIB does not exhibit over-integration on this compositionally imbalanced data.

These results demonstrate that the probabilistic and uncertainty-aware design in multiVIB provides an advantage in avoiding the hidden failure modes of both shared feature- and paired cell-based integration, enabling more reliable and biologically coherent alignment even under deliberately adversarial conditions. We next quantitatively evaluate the performance of multiVIB in different integration scenarios against state-of-the-art tools, highlighting its particular advantage as a unified framework for mosaic datasets.

### 3.2 multiVIB delivers robust, biologically coherent integration across diverse scenarios

Building a multimodal cell atlas from collaborative efforts requires integrating mosaic datasets where some modalities are anchored through jointly profiled cells while others can only be anchored with shared features. It is therefore critical that the integration method performs reliably well in both horizontal (feature-shared) and vertical (joint-profiled) settings. Existing tools often excel in one category or the other but rarely both, forcing researchers to set priorities and selectively sacrifice between batch correction, modality alignment, and biological fidelity when assembling custom hybrid workflows.

multiVIB was designed to mitigate this trade-off: across different integration settings, its unified model backbone should perform equally well to preserve cell-type structure while aligning modalities and removing study-specific variation. To assess whether multiVIB fulfills this promise, we conducted a systematic benchmarking analysis against 11 state-of-the-art approaches, using the jointly profiled 10X Genomics Multiome data, and 4 additional single-modality datasets from independent studies using human PBMC and bone marrow samples[38–41].

We evaluated the integration performance with quantitative metrics in the following aspects[14]: (1) *batch correction* within each modality, measured by batch mixing score; (2) *modality alignment* across modalities, measured by modality mixing score and average silhouette width (ASW) modality; (3) *biological cell-type preservation*, assessed by ASW cell type at both coarse (L1) and granular (L2) label resolutions; and (4) *cross-modal mapping*, measured by overall accuracy and macro F1 score through label transfer task. Details of these metrics are described in *Methods*. For vertical integration, we compared multiVIB against 4 leading methods: multiVI[27], SMILE[26], Cobolt[25], and Seurat (dictionary learning)[28] (Figure 2D). multiVIB showed superior performance in modality alignment and cross-modal mapping, while all methods achieved comparable scores in cell-type preservation. In addition, multiVIB and Seurat (dictionary learning) outperformed other methods in batch correction for both modalities, underscoring their broader utility. For horizontal integration that uses shared features between these two modalities as anchors, we converted scATAC-seq peak matrix into gene activity matrix to match with gene expression features of scRNA-seq data, following a well-established practice in horizontal integrations[19, 20]. We compared multiVIB against 7 state-of-the-art methods: Seurat (CCA)[19], Scanorama[10] (RRID:SCR_021539), BBKNN[12] (RRID:SCR_022807), Harmony[11] (RRID:SCR_022206), scVI[8] (RRID:SCR 026673), sysVI[42], and LIGER[20] (RRID:SCR_018100) (Fig. 2E). multiVIB demonstrated consistently strong performance in all four benchmarking categories, similar to scVI and Seurat (CCA). Notably, multiVIB excelled simultaneously in modality alignment and batch correction (Fig. 2E). Particularly, we examined the accuracy of label transfer per cell type, and we observed that multiVIB is the leading method in both vertical and horizontal integration that has a balanced F1 score per cell type (Supplementary Fig. S2 and S3). This further demonstrates the advantage of integration through multiVIB in the task of cross-modality label transfer.

To quantify the balanced performance of multiVIB across tasks, we compared all methods by ranking the average scores across all four benchmarking metrics[14] (Fig. 2D-E). We found that horizontal integration tools (e.g., Harmony, Scanorama, and BBKNN) perform well on cross-studies harmonization but falter on cross-modal alignment; and vertical methods (e.g., Cobolt, SMILE, MultiVI) excel in vertical settings for which they were designed, but struggle to generalize across studies. In contrast, multiVIB maintained top performance across all categories simultaneously. This consistently strong performance is what enables multiVIB to function as a unified framework for mosaic data integration without manual tuning of integration strategies or custom assembly of workflows.

We further validated whether the unified latent space produced by multiVIB supports biologically meaningful downstream tasks in the multimodal, multi-study integration, specifically label transfer and cell querying. Label transfer tests the alignment across modalities at the cell type level. Using Granja *et al*. RNA-seq data[39] as the reference and Satpathy *et al*. ATAC-seq[40] as the query, multiVIB transferred cell-type annotations with high concordance to the original biological labels from the independent study[40], (normalized mutual information or NMI = 0.616 and 0.578 for coarse (L1) and detailed (L2) cell-type annotation, respectively, Supplementary Fig. S4B, C). This result demonstrates reliable cross-modal mapping despite substantial differences in data modality and experimental conditions. Cell querying across datasets evaluates whether the unified latent space preserves biologically meaningful neighborhood structure. Again, treating Satpathy *et al*. ATAC-seq data[40] as queries, we identified their 15 nearest neighbors in both 10X[37] and Granja *et al*.[39] RNA-seq datasets. Retrieved neighbors from both studies consistently matched to the query cell types (Supplementary Fig. S4D), demonstrating that multiVIB maintained coherent cell-type structure across datasets and modalities. These results confirmed that multiVIB not only performs well on benchmark metrics but also delivers reliable, biologically grounded alignment that supports real analytical tasks.

Furthermore, we also benchmarked the batch correction task in human lung cell atlas (scRNA-seq)[43] and mouse brain cell atlas (scATAC-seq)[14]. To evaluate methods from the aspects of biological conservation and batch correction, we followed the benchmark study by Luecken *et al*. and primarily selected metrics that evaluate biological conservation and batch correction in latent space[14]. In this benchmarking, we also included macro F1 and overall accuracy to evaluate methods in the task of cross-dataset label transfer. We compared multiVIB against 7 state-of-the-art methods: Seurat (CCA)[19], Scanorama[10], BBKNN[12], Harmony[11], scVI[8], sysVI[42], and LIGER[20]. multiVIB stood out as one of the top methods for human lung cell atlas scRNA-seq integration (Supplementary Fig. SSA). For scATAC-seq datasets (mouse brain cell atlas), while most methods performed poorly for batch correction[14], multiVIB and scVI led over other 6 methods and showed clear advantage in integration of scATAC-seq (Supplementary Fig. SSB). Luecken *et al*. pointed out scANVI[44] as a leading method that consistently integrate datasets well for both scRNA-seq and scATAC-seq[14]. However, scANVI stands out from all these horizontal methods because it uses partial cell-type labels to perform integration. This inherently gives scANVI an advantage to learn integrated latent space that better preserves biological information and serves cross-data label transfer. For a fair comparison, we incorporated classification loss in training multiVIB and compared with scANVI in both human lung cell atlas and mouse brain cell atlas data. We noticed that both multiVIB and scANVI, trained with partial cell type labels, showed improved performance from multiVIB and scVI that were trained without any supervision (Supplementary Fig. SSC). Meanwhile, multiVIB with partial labels slightly outperformed scANVI in human lung cell atlas (scRNA-seq) and mouse brain cell atlas (scATAC-seq).

### 3.3 Scalable and generalizable mosaic integration of single-cell multimodal data from mouse primary motor cortex

The mouse primary motor cortex (MOp) was the central focus of the first stage of BRAIN Initiative Cell Census Network (BICCN)[45]. As a result, mouse MOp has become one of the brain regions that has been most comprehensively profiled. While the reference atlas has been established using deeply standardized experimental and computational pipelines[46], other research groups have studied the same brain region using additional molecular assays, offering complementary perspectives on neuronal identity and cellular state. Integrating these diverse datasets is key to constructing the next-generation cell atlas that reflects the full molecular complexity of the brain. Here, we assembled a compendium of 9 modalities from 5 additional published sources[23, 24, 47– 49], alongside the BICCN reference atlas[46] to create a testbed for evaluating how multiVIB harmonizes diverse omics layers into a coherent cell atlas (Fig. 3A).

**Figure 3:**
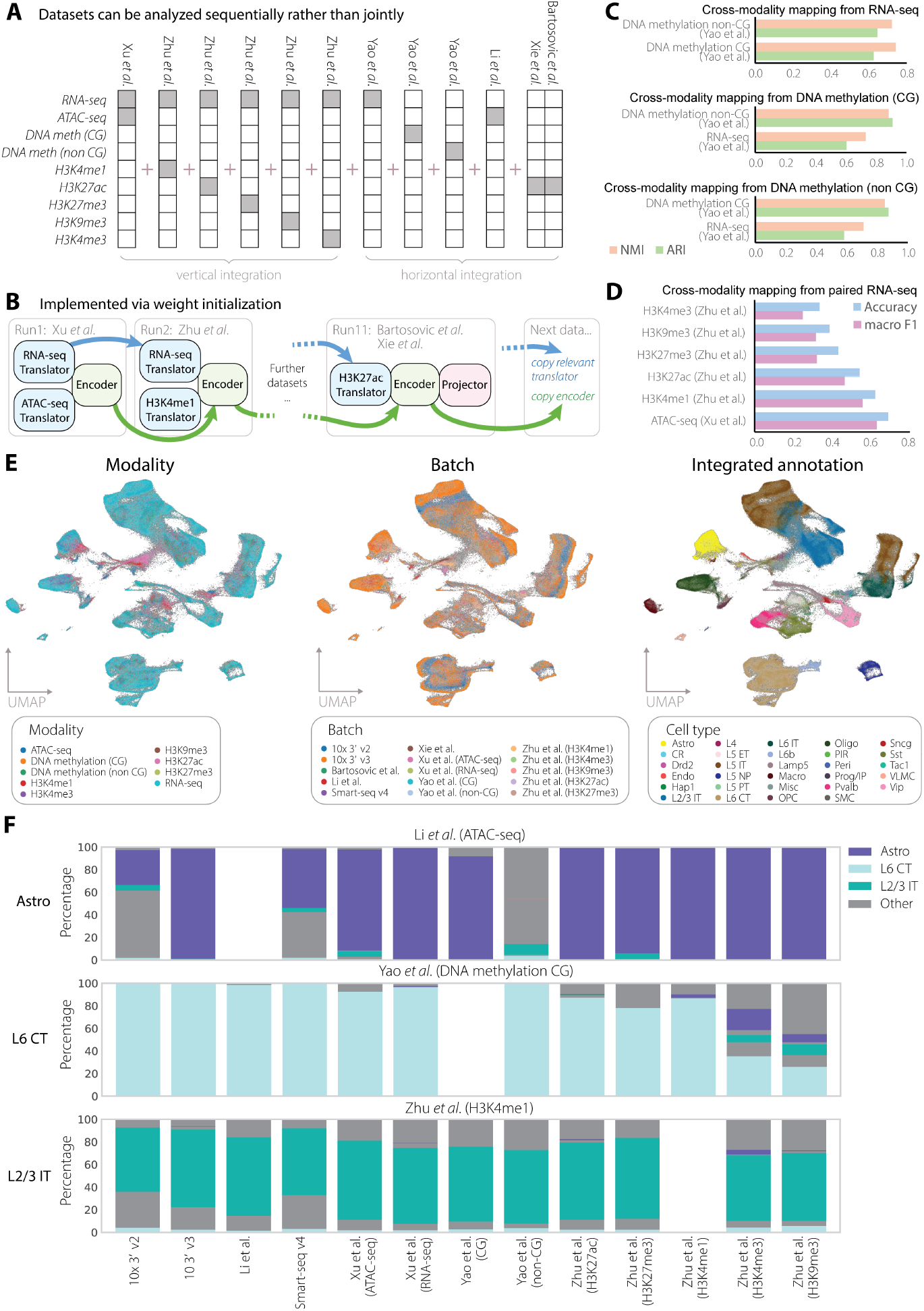
Mosaic integration of mouse motor cortex data over 9 modalities. (A) Overview of MOp data included in this mosaic integration task. Datasets were collected from 6 sources covering 9 modalities. multiVIB was performed in vertical integration mode on data with joint measurements, such as RNA-seq with ATAC-seq or histone modifications. And for data without joint measurements, such as DNA methylation, the horizontal integration mode of multiVIB was implemented. (B) Dataset-by-dataset training strategy for integrating MOp data. The encoder always inherits weights from previous training. Translators are updated from previously relevant translators or created from scratch. (C-D) Evaluation of integration with cross-modal mapping, also known as label transfer. (C) Evaluation of horizontal integration. ARI and NMI metrics were used to measure label transfer from one modality to the others. (D) Evaluation of vertical integration. Macro F1 and overall accuracy scores of 6 epigenomic modalities are computed via label transfer from their paired RNA-seq data. (E) Visualization of mouse motor cortex data via UMAP. Cells are colored according to modality sources (left), batch identities (middle), and unified cell type annotations (labels transferred from Yao *et al*. to all cells, right). (F) Cross-study and -modality cell querying. Astrocytes, L6 CT cells, and L2/3 IT cells from 3 different studies (the panels) were queried against all integrated cells to identify the most similar cells in all other data sources (the x-axis labels). Each bar represents the total cell population from each study, color coded by the identified cell types. Aside from the 3 queried cell types, all other cell types are colored gray. These cell types and source datasets were selected because they are well represented across all datasets in the integration and yield high-confidence label transfer, making them suitable for illustrating embedding consistency. Modalities with lower label transfer accuracy (e.g., H3K4me3 and H3K9me3; panel D) are not shown as their inclusion would reflect label transfer failure rather than embedding structure.

In addition to the mosaic data scenario, this integration task presents several common challenges: (1) the dataset composition was heavily skewed toward single-modality scRNA-seq, which is a common imbalance in large single-cell compendium; (2) although jointly profiled data provided direct RNA–ATAC correspondences as cell anchors, the vast majority of ATAC-seq cells were profiled independently without paired RNA measurements; and (3) no jointly profiled data linked gene expression and DNA methylation, restricting those modalities to horizontal integration based solely on shared genomic features. Moreover, training all these nine modalities at once is computationally expensive and would prevent more incoming data. To address these practical and conceptual challenges, we implemented multiVIB with its continuous dataset-by-dataset learning strategy: the shared encoder always inherited weights from prior training, while translators were either initialized from previously learned components or trained *de novo* when encountering new modalities (Fig. 3B). This strategy underscored multiVIB’s flexibility to combine vertical and horizontal integration within a unified probabilistic framework, which enables large-scale singlecell atlases to accumulate datasets, including more new modalities in an evolving fashion without retraining from scratch. Note that training order had negligible impact on the quality of this particular data integration task because all major cell types were represented across datasets. As a result, the shared latent space learned from earlier datasets remained a suitable scaffold for subsequent additions. But if a newly introduced dataset contains cell types absent from all previously seen datasets, the latent space may shift to accommodate this new variation at the expense of earlier representations. We therefore recommend starting sequential training from the dataset with the broadest cell-type coverage.

To validate the biological information in the learned cell embeddings, we evaluated cross-modal mapping of how accurately cell-type annotations could be transferred across datasets and modalities. For horizontal integration tasks, multiVIB achieved high label concordance across 3 modalities (RNA, DNA methylation in CG and non-CG contexts). In particular, both ARI and NMI reached nearly 0.9 in the mutual transfer between CG and non-CG DNA methylation data (Fig. 3C), indicating effective alignment of closely related molecular readouts. For vertical integration tasks, multiVIB achieved label transfer accuracy of 0.63 macro F1 and 0.69 overall accuracy for ATAC-seq data (Fig. 3D). However, the accuracy decreases to 0.23 (macro F1) and 0.32 (overall accuracy) in H3K4me3, potentially reflecting the greater biological distance between transcriptional output and distal epigenetic regulation. UMAP visualization further supported these trends, showing that multiVIB yielded well-mixed cell embeddings across studies and modalities while preserving coherent cell-type structure (Fig. 3E). Together, these results demonstrated that multiVIB reconstructs biologically meaningful relationships across heterogeneous molecular layers and achieved a strong alignment when modalities share regulatory context.

Finally, we show that the unified cell embedding learned by multiVIB empowers biologically meaningful cross-source and cross-modal cell querying. Using representative cell types (astrocytes, L6 CT, and L2/3 IT cells) as queries, we searched for their most similar counterparts across multiple datasets. Query searching consistently retrieved cells with matched identity across many modalities and data sources, demonstrating that the integrated embedding preserved biological correspondence beyond simple alignment (Fig. 3F). However, the agreement decreased in histone modification modalities such as H3K4me3 and H3K9me3, consistent with their lower label transfer performance. Meanwhile, we noticed that CG and non-CG feature representations are highly correlated at the population level (Figure 3C), yet label transfer accuracy is notably higher for CG-derived annotations for the Li et al. ATAC-seq dataset. This discrepancy arises because the two separated regions, either CG or non-CG regions, on the genome may resolve cell-type boundaries at different granularities, causing cells near cluster boundaries to be assigned to different types depending on which feature representation is used. This highlights a general limitation: global correlation between feature representations does not guarantee consistent label transfer accuracy across fine-grained cell-type boundaries. These results suggested that these layers may capture complementary rather than directly mappable aspects of cell identity (see Discussion).

### 3.4 Disentangling conserved and species-specific molecular programs across primate basal ganglia

The basal ganglia are the primary locus of pathology in motor and cognitive disorders ranging from Parkinson’s to addiction[50–52], making the validation of non-human primate (NHP) models a critical priority for translational neuroscience[6]. While the anatomical organization of basal ganglia is broadly conserved[53], the extent to which molecular programs are shared across human and NHP remains an open question[54]. To establish this translational baseline, the BICAN consortium has generated high-resolution single-cell transcriptomic profiles across human, rhesus macaque, and common marmoset (Johansen *et al*.[3]). Using this dataset, we validated the application of multiVIB in comparative analysis by constructing a unified cross-species landscape and recovering the consensus taxonomy. We further investigated the species-specific embeddings learned by multiVIB to recover distinct transcriptional programs that characterize the human striatum.

We first demonstrate the capacity of multiVIB in the task of cross-species integration with primary motor cortex data from human[55] and mouse[46]. Similar to batch correction benchmarking, we evaluated performance of 8 horizontal methods over two main categories, biological conservation and species alignment respectively. Even though biological conservation score shows most methods, except BBKNN and LIGER, succeeded in preserving biological information, the majority failed to mix the two species (Supplementary Fig. SS). We observed that multiVIB, alongside sysVI and Seurat (CCA), are the only methods that can clearly align cells across species, as indicated by higher species alignment scores (Supplementary Fig. SS). With this support of cross-species integration benchmarking, we next deployed multiVIB to carry out integration of transcriptomics data across species under the horizontal principle, utilizing the two-phase training strategy (Fig. 1F), with homologous genes as biologically grounded anchors (Johansen *et al*.[3]). In the first phase, the model focused on cross-species alignment by constraining the mapping within shared features across species only. The mask in the linear translator prevented spurious correlations between genes across species, ensuring that the early goal of model training was cross-species alignment (Fig. 4A). In the second phase, we relaxed this constraint and allowed the model to recover species-specific structure in the data (Fig. 4B).

**Figure 4:**
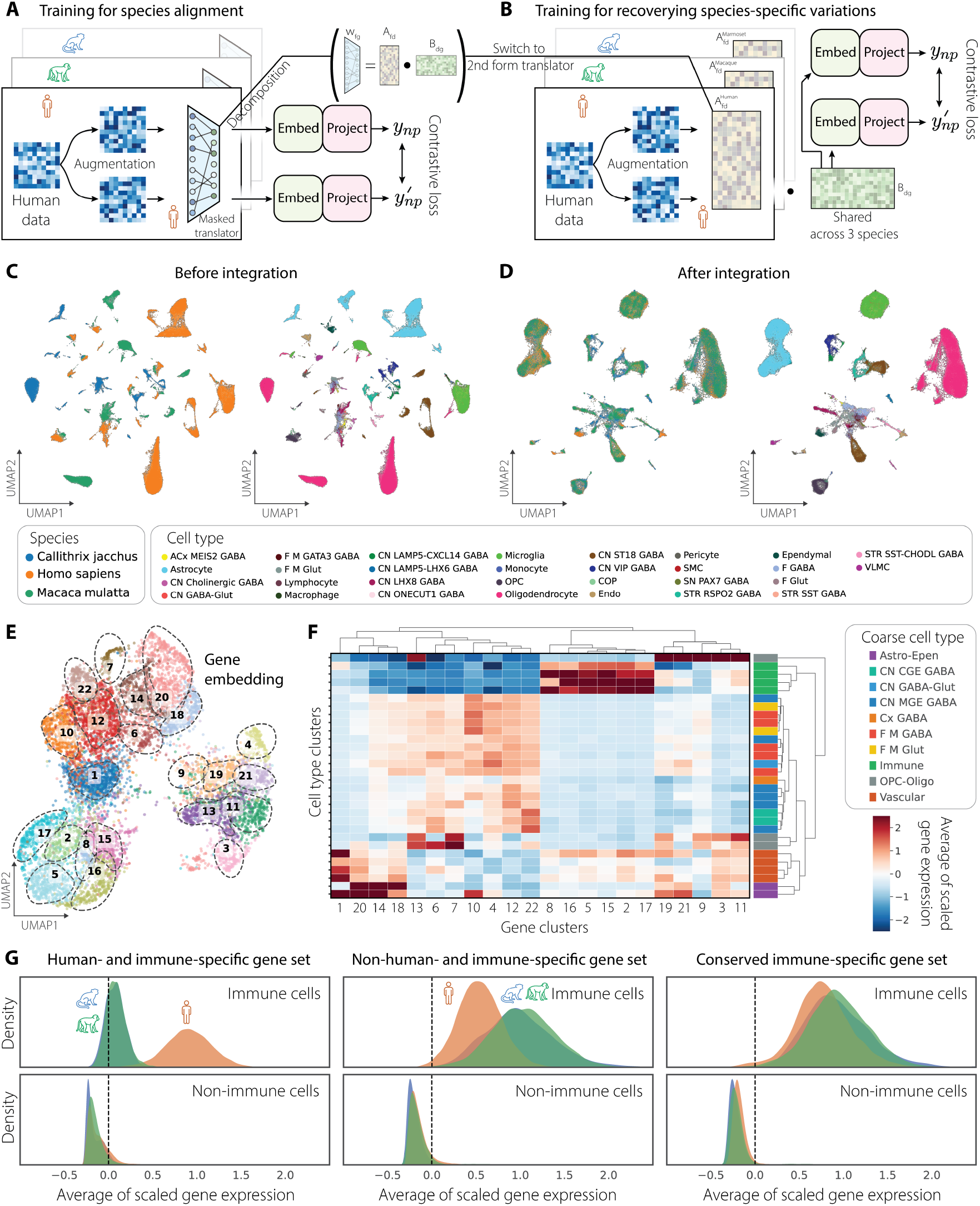
Horizontal integration of basal ganglia data across 3 species. (A) Phase 1 training focuses on species alignment using a masked linear translator to restrict mappings between non-homologous and conserved homologous genes. (B) Phase 2 training recovers additional species-specific variations. The trained translators from phase 1 are decomposed into two lower-rank components, *A*_*fd*_ and *B*_*dg*_, which are updated without any masking. (C) UMAP visualization of cell embedding before integration. Cells are colored by species and cell type. (D) UMAP visualization of cell embedding after multiVIB integration. (E) Co-embedding of genes from all 3 species. All genes from 3 species are co-embedded through the learned *A*_*fd*_ matrices. Then, louvain clustering is applied to group all genes into distinct clusters. (F) Scaled average expression of gene clusters in cell types, illustrating cell-type-specific enrichment. (G) Probability density of averaged scaled gene expression plotted for immune-related gene programs, showing recovery of (1) human-specific, (2) non-human primate-specific, and (3) conserved immune gene programs, obtained from the learned translator matrices after integration. Y-axes are individually scaled, but the area under each curve integrates to 1.

First, we confirmed that multiVIB effectively aligned human, macaque, and marmoset cells into a unified latent space, grouping transcriptomic profiles by homologous cell types rather than species (Fig. 4C-D). This alignment was consistent with the consensus taxonomy defined by the transcriptomic atlas (Johansen *et al*.[3]), with major cell classes and subclass boundaries preserved across species. Using the harmonized cell type annotation from Johansen *et al*.[3] as the ground truth, we benchmarked multiVIB against other 7 horizontal methods, through evaluating biological conservation and species alignment (Supplementary Fig. SS). multiVIB demonstrated consistently high quality of cross-species integration as a leading method (Supplementary Fig. SS). Beyond reliably aligning cells of 3 species, multiVIB also presents the advantage of identifying species-specific regulatory factors. Next, clustering of the learned gene embedding (Fig. 4E) revealed gene groups with strong cell-type-specific enrichment (Fig. 4F), indicating that multiVIB captured coordinated gene modules that underlie the cellular identity. To further assess multiVIB’s capability of preserving species-specific signals, we examined immune-related gene modules as a representative case (Fig. 4G). Analyzing species-specific matrices learned during training, we recovered three biologically distinct categories of immune gene sets: human-specific, NHP–specific, and conserved across all three species (details in *Methods*). Successful retrieval of coherent, species-resolved gene programs demonstrated that multiVIB not only aligned shared cell identities but also preserved evolutionarily meaningful divergence.

## 4 Discussion

A central goal of single-cell biology is to construct multimodal atlases that capture the full diversity of cellular states across molecular layers. The BRAIN Initiative has been advancing this vision with rapid generation of large-scale transcriptomic and epigenomic datasets[56]. With reference atlases developed by individual centers and datasets analyzed within specific scope of research interest, integrating these disparate efforts into a unified knowledge resource remains a major challenge. Existing computational methods are typically designed for limited integration scenarios, underscoring the need for a more versatile and generalizable framework for single-cell multi-omics data integration. multiVIB fills this critical gap as a unified framework for integrating increasingly heterogeneous datasets while preserving biological fidelity. By coupling a minimalist model architecture with a probabilistic contrastive learning strategy, we show that multiVIB adapts to diverse integration scenarios and mitigates artifacts such as spurious alignments that frequently arise in multimodal data integration. In this study, we also demonstrated that this capability enables reference atlases to evolve continuously as new datasets accumulate (Fig. 3) and supports cross-species integration for comparative analyses of brain cell types (Fig. 4).

### 4.1 Principles for atlas-scale single-cell integration

Our experience applying multiVIB to more than one million cells across BICCN and BICAN datasets revealed several general principles that extend beyond any individual method. These principles reflect recurring patterns in multimodal or multi-species atlas construction and help explain why naive or purely technical integration strategies often fail to preserve true biological variation.

First, data representativeness is critical. Jointly profiled multi-omics datasets provide powerful anchors for vertical integration, yet they remain sparse relative to the scale of single-modality datasets and often undersample rare or transitional cell populations. Over-reliance on these incomplete anchors can propagate sampling biases, leading to spurious alignments or the collapse of biologically meaningful states. Second, integration should preserve biological interpretability.

Attempts to force heterogeneous assays into superficially shared feature spaces—for example, summarizing chromatin conformation or accessibility peaks as “gene activity” proxies—can distort the underlying regulatory logic. Such engineered features could assist data alignment across modalities but may frequently obscure modality-specific information rather than enabling genuine integration. Third, multimodal integration requires semantic awareness. Modalities that appear structurally similar may encode opposing regulatory states. For instance, chromatin accessibility and DNA methylation can reference the same genomic regions yet convey inverted functional meanings. Treating these modalities as interchangeable features risks misinterpreting the biological signals required to distinguish conserved programs from species- or lineage-specific regulation. To guide the selection and development of integration methods in line with these principles, we summarized our recommendations in Table 1 based on the benchmarking results in this study.

**Table 1:** Comparison of representative integration tools across four scenarios. Both 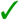 and 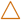 indicate supported integration tasks, with 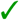 being our recommendation. * indicates that the method was not tested in this study.

| Tool | scRNA-seq<br>Batch | scATAC-seq<br>Batch | Cross-Modal | Cross-Species |
| --- | --- | --- | --- | --- |
| multiVIB | ✓ | ✓ | ✓ | ✓ |
| scVI | ✓ | ✓ | ✓ | ✓ |
| sysVI | ✓ | ✗ | ✗ | ✓ |
| multiVI | * | * | ✓ | ✗ |
| BBKNN | ✓ | △ | ✗ | ✗ |
| Cobolt | * | * | △ | ✗ |
| Harmony | ✓ | △ | △ | △ |
| LIGER | ✓ | △ | △ | △ |
| Scanorama | ✓ | ✗ | ✗ | △ |
| Seurat | ✓ | △ | ✓ | △ |
| SMILE | * | * | △ | ✗ |

Together, these observations highlight that multimodal integration is not merely a problem of statistical harmonization but fundamentally one of biological inference. Preserving the diversity of regulatory mechanisms across assays, species, and modalities is essential for reconstructing cellular identity at atlas scale. These principles clarify a broader role of multiVIB within the BRAIN Initiative. By explicitly modeling uncertainty, multiVIB balances alignment with biological preservation, avoiding both under-integration and over-correction while maintaining the semantic structure of each modality. The modular design of multiVIB also enables scalable integration across the rapidly expanding scope of BICAN. The challenges encountered in integrating BICCN multimodal motor cortex datasets and BICAN cross-species basal ganglia datasets will only get intensified as additional modalities, developmental stages, and disease contexts are incorporated. In this context, multiVIB provides a generalizable computational foundation for constructing the next generation of comprehensive multimodal brain cell atlases.

### 4.2 Addressing spurious alignment in single-cell data integration

Spurious alignment remains a persistent challenge in single-cell data integration, arising when models converge on solutions that minimize technical variation rather than capturing true biological correspondence. In horizontal integration, this often stems from poorly engineered cross-modal shared features; in vertical integration, the risk increases when paired cells fail to represent the full spectrum of cellular diversity. multiVIB mitigates spurious alignment in both scenarios through its two-phase modality-specific translator design for horizontal integration and probabilistic contrastive embeddings for vertical integration. Compared to reconstruction-based approaches, contrastive loss provides a more well-defined alignment objective that explicitly strengthens biological correspondence across datasets, as supported by our conceptual stress test showing improved cross-modal alignment of multiVIB over multiVI (Figure 2B and C). Although both multiVIB and scVI are grounded in variational inference, they differ fundamentally in their learning objectives: reconstruction loss in scVI drives the latent space to capture cell states, whereas contrastive loss in multiVIB directly optimizes for biologically meaningful cross-dataset alignment. We argue that these two mechanisms shape the latent space in intrinsically different ways, even when they yield similar performance under standard evaluation metrics, and that understanding this distinction is important for designing more reliable integration methods.

Although a detailed investigation is beyond the scope of this study, it would be valuable to further explore how reconstruction and contrastive objectives shape a model’s ability to approximate biological variation, particularly in representing cell states in latent space. Jointly optimizing these objectives within a unified integration framework may enhance compositional representation learning, but such effort should be grounded in a deeper understanding of their respective roles in representation learning for single-cell data. Notably, a recent study by Bahrami *et al*. highlights how contrastive pretraining could reshape current single-cell foundation models beyond traditional reconstruction-based objectives, suggesting that contrastive learning may offer a new perspective for understanding cell states in latent space[57].

### 4.3 Supporting ontology-level cell type harmonization

A major objective of many consortium projects, including the BRAIN Initiative, is to establish unified and interoperable cell-type taxonomies across datasets, modalities, and species[1]. Despite ongoing community efforts to standardize nomenclature and ontology, practical approaches for data harmonization remain challenging, as each study often defines cell populations at different resolutions or through distinct molecular modalities. This challenge has motivated the development of automated tools such as CellHint[58], which performed cross-dataset label harmonization by identifying equivalent cell populations in the shared cell embedding. Our results showed that the unified latent space learned by multiVIB provides a strong computational foundation for ontology-level harmonization. When CellHint was applied to multiVIB-derived embeddings, it achieved high cross-study concordance across transcriptomic, chromatin accessibility, DNA methylation, and histone modification datasets, indicating that the learned representations captured biologically coherent cell relationships that generalized effectively to external harmonization frameworks (Supplementary Fig. S8). This capability positions multiVIB not only as a method for data integration but also as an enabling infrastructure for the BRAIN Initiative’s broader goal of building interoperable, cross-consortium cell atlases grounded in a unified cell ontology.

In this study, we also briefly examined how incorporating partial cell type labels can enhance the integration performance of multiVIB. Providing the model with available cell type annotations can help reduce the risk of cell state collapse and spurious alignment. However, most existing approaches treat cell type information as flat labels. While such supervision improves the model’s ability to learn discriminative latent representations across cell types, it may obscure more nuanced relationships among cell states, particularly in cellular systems that contain hierarchical or transitional structures among cell states. Therefore, unsupervised integration methods remain valuable for faithfully capturing the continuous and structured relationships among cell states. A key challenge, particularly in large-scale efforts such as BICAN, is how to establish ontological cell type annotations through unsupervised learning. We envision that multiVIB could support ontology-level cell type harmonization, where integration and annotation are iteratively refined through a feedback loop that incorporates ontology-structured supervision.

### 4.4 Limitation

In this study, we demonstrated that contrastive learning is effective in bridging different modalities into a unified latent space. However, this approach carries the risk of erasing modality-specific variation in the process of identifying a shared representation. As more modalities are incorporated, the shared intersection may become increasingly constrained. By additionally encouraging the preservation of variation within each dataset, multiVIB mitigates this effect and reduces the risk of collapsing cell states across modalities into an overly restrictive common space. While learning a unified cell embedding across studies, species, and modalities remains an appealing goal for constructing comprehensive cell atlases, our results suggest that such a fully unified representation may not always be biologically attainable. For example, we noticed some major cell types were poorly separated using histone modification features such as H3K9me3 and H3K4me3. Although this could reflect technical limitations, such as insufficient measurement or experimental sensitivity, it is also possible that some epigenetic marks simply cannot encode cell identity at the same level of specificity as transcriptional or chromatin accessibility signals. Consistent with this interpretation, we observed that current computational methods, including multiVIB, performed worse in cross-modal label transfer compared to within-modality label transfer, which could reflect fundamental biological divergence across omics layers besides technical limitations. These findings suggest that future efforts on the development of computational methods should prioritize extracting modality-specific biology instead of imposing a universal embedding for all modalities. Methods that can better capture the complementary roles of distinct molecular modalities in defining cellular state and function may result in a more faithful and multidimensional representation of cell identities.

## 5 Methods

### 5.1 Probabilistic contrastive learning

We approximate the latent representation using an encoder network that produces a posterior distribution in the latent space, and we train the model using a contrastive loss. These elements form the core of probabilistic contrastive learning in our study.

The conceptual framework for the loss function in multiVIB has roots in the information bottleneck (IB) study[59]. The objective of IB is to maximize the mutual information contained in latent representation *z* from input data *x*, while restricting how much information about the identity of each data element is allowed in this representation. Under the information bottleneck principle, a variational form of IB (VIB)[33] can be set up as:

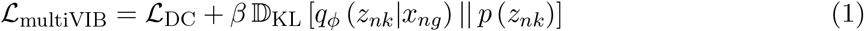

where ℒ_DC_ is our task-specific contrastive loss and *q*_*ϕ*_ (*z*_*nk*_|*x*_*ng*_) is the variational posterior distribution for the latent variables conditioned on the data, implemented as an encoder network, and *p* (*z*_*nk*_) is the prior over latent representation. 𝔻_KL_[·||·] represents the KL divergence. The hyperparameter *β* sets the bottleneck strength, which controls the amount of information retained in *z*_*nk*_. In the Eq. (1), the KL divergence term ensures this distribution does not stray too far from the prior. In this study, we set *β* as 0.05 for all data integration.

Contrastive learning has been successfully applied to learn representations of images in numerous studies[60, 61]. This was soon copied to other data types, including single-cell omics data[26, 62–64]. The key component of contrastive learning is to use data augmentation to create positive pairs that represent different “views” of the same input data. These positive pairs, along with negative examples, are combined to compute a contrastive loss, which maximizes embedding similarity between positive pairs while maximizing differences among negative examples. We use the “decoupled contrastive learning” (DC) loss from Yeh *et al*.[65].

During horizontal integration, we apply this contrastive loss to the projector outputs *y*_*np*_, as written above (Eq. 4). The (*v*) superscript refers to two augmented views of the same data point. The loss terms associated with cell *n* would be written:

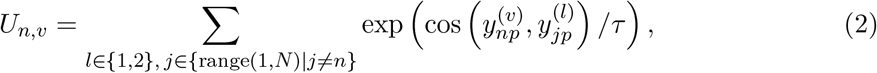

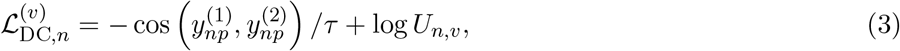

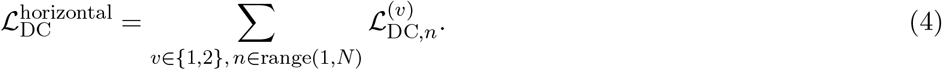

*U*_*n,v*_ is a latent space distance metric summed over all pairs of negative examples for the view *v* of cell *n*. The sum is composed of 2(*N* − 1) terms, since each example has two augmented views, where *N* is the number of cells in a minibatch.

During vertical integration, we use the latent space *z* to compute the loss instead. The full loss for vertical data integration (again for the case of two modalities A and B) is given in Eq. 7. In the vertical case, the (*v*) superscript refers to measured data from different modalities rather than data views created by augmentation. For example if we are choosing two modalities (A and B) as our positive examples (different views of the same data), the loss terms associated with cell *n* would be written

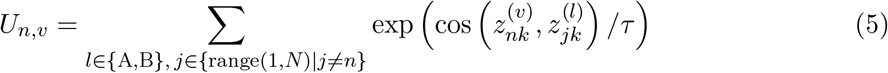

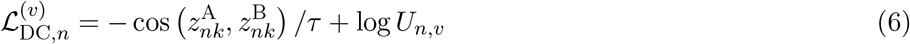

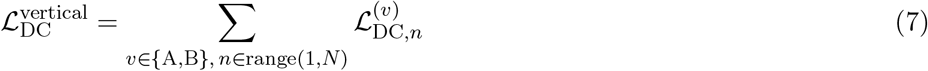

Of note, the two contrastive loss terms 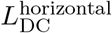 and 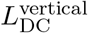 serve distinct but complementary roles in our framework. The horizontal contrastive loss, 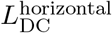, is applied to multiple views of the same modality and is designed to capture and preserve intrinsic variation within that modality. In contrast, the vertical contrastive loss, 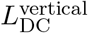, operates across modalities for the same cell, encouraging alignment of modality-specific representations within a shared latent space.

This separation of objectives allows the model to balance two competing goals: maintaining modality-specific structure while achieving cross-modal integration. The joint optimization of these two loss terms therefore provides a mechanism to regulate this trade-off, enabling the model to learn representations that are both biologically informative within each modality and coherent across modalities.

The probabilistic nature of multiVIB stems from its variational encoder, which defines a distribution over the latent space rather than a single deterministic embedding. Representing cells as probability distributions adds quantifiable uncertainty to cell embeddings[34, 35], which we argue is essential for capturing the variable and potentially limited information contributed by each data modality.

To motivate this design choice, consider a scenario in which a new data modality reveals a previously uncharacterized T cell variant absent from all other modalities in the training data. Because the model has no direct supervision for this cell state across modalities, a deterministic encoder would produce an arbitrary or poorly constrained embedding. A probabilistic encoder, by contrast, can infer the embedding of this unobserved cell state through its distributional proximity to related T cell variants that *are* observed in other modalities. Crucially, this uncertainty-aware inference also suppresses spurious cross-modal alignment: when evidence from a modality is weak or absent, the broader posterior naturally limits overconfident integration.

However, the strength of the imposed prior introduces a practical trade-off. A stronger KL divergence penalty encourages greater mixing across datasets but can erode the discriminative power needed to resolve distinct cell types. To characterize this trade-off empirically, we examined the effect of the KL divergence weight on two example datasets. Our experiments indicate that a weight in the range of 0.02–0.05 achieves a balance between biological variation preservation and cross-modal alignment when combined with contrastive learning (Supplementary Fig. S9).

### 5.2 The backbone of multiVIB

The model backbone of multiVIB consists of three parts: (1) a modality-specific linear translator, a shared encoder, and (3) a shared projector.

#### Notation

Let 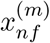 denote the observed feature vector for cell *n* in modality *m*, where *f* indexes modality-specific features. Let *g* index genes in the shared gene-centric feature space, *k* the dimensionality of the shared latent cell embedding, and *p > k* the dimensionality of the final projection space. Covariates such as batch, species, or modality are denoted *C*_*n*_. The backbone of multiVIB consists of three mappings:

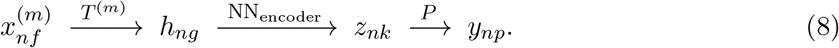

Each modality-specific translator *T* ^(*m*)^ is implemented as a linear map with weights *W* ^(*m*)^ ∈ ℝ^*f ×g*^ and bias 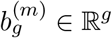.

#### Modality-specific translator

Each modality *m* is equipped with a linear translator mapping modality-specific features into the shared gene expression space:

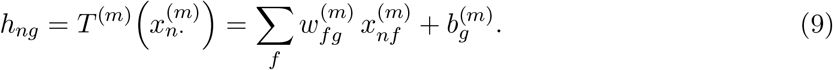

The gene expression-centered translated space *h*_*ng*_ leverages the abundance of scRNA-seq datasets while providing an interpretable coordinate system. The linear structure of *T* ^(*m*)^ makes 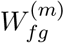 and 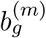 directly interpretable for uncovering cross-modal feature relationships and regulatory interactions.

We implemented two different forms of translators for distinct purposes. The first form of the modality-specific translator is implemented as a fully-connected dense layer. The original input 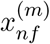 is multiplied by 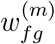 and summed with a bias term 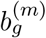, giving the translated output 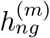, where *f* stands for the original feature space and *g* is the shared feature dimension. However, this dense layer could learn implausible correlations between features *f* and *g*, which we believe would be a major source of spurious alignment. Therefore we add a binary mask matrix, *m*_*fg*_ ∈ {0, 1}, informed by a biological prior, to force the model to explore only “biologically-plausible” correlations between the original feature space *f* and shared feature space *g* (Eq. (10)).

Such a biological prior can come in the form of two features that are physically close in the genome. For example, opening of a promoter region in ATAC-seq data is more likely to influence transcription of the proximal gene than a distant gene. 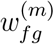 is sparse, due to the sparsity imposed by the mask. Meanwhile, imposing the mask also prevents the translator from recovering dataset-specific variations. Therefore, we introduce the second form of the translator through lower-rank matrix decomposition (Eq. (11)). The masked weight matrix *w*_*fg*_*m*_*fg*_ is decomposed into 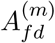 and 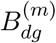 using singular value decomposition (Eq. (12)). *d* is a compressed dimension (128 in multiVIB), making this scheme equivalent to allowing only low-rank updates to the weight matrix. Without the mask, the second form of the modality-specific translator gives the model more flexibility to explore feature correlations between the original and shared feature space. In the training detail, we will further explain how we interchange these two forms to serve integration purpose.

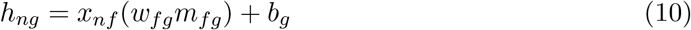

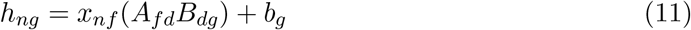

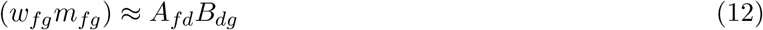

#### Shared probabilistic encoder

The shared encoder, denoted NN_encoder_, produces the parameters of a Gaussian distribution rather than a single deterministic embedding. Specifically, it outputs a cell-specific mean and log-scale of a Gaussian posterior distribution:

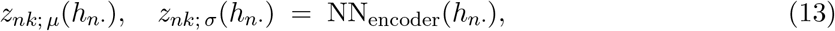

where *σ* denotes the (diagonal) standard deviation. A stochastic latent representation *z*_*nk*_ is then obtained via the reparameterization trick. This probabilistic formulation, along with a standard normal prior imposed on *z*_*nk*_ imposes an information bottleneck analogous to VAEs, stabilizes training, and regularizes the latent biological state. The resulting *z*_*nk*_ serves as a modality-agnostic embedding across datasets.

*z*_*nk*_ is a *K*-dimensional latent variable representing the “unified” cell embedding, with a standard normal prior (Eq. 14). The encoder NN_encoder_ is implemented as a multi-layer perceptron. Here, we use a fully-connected dense neural network with 3 hidden layers. The encoder generates sufficient statistics for the posterior distribution *q*_*ϕ*_(*z*_*nk*_ | *x*_*ng*_*′*), namely *z*_*nk*; *µ*_[*x*_*ng*_*′*] and *z*_*nk*; *σ*_[*x*_*ng*_*′*] (Eq. 15). The posterior *q*_*ϕ*_(*z*_*nk*_ | *x*_*ng*_*′*) follows a normal distribution (Eq. 16). Here *ϕ* represents the bundle of parameters in the translator and encoder. We update these model parameters using variational inference[33, 66].

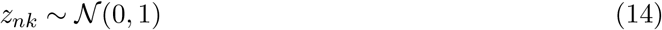

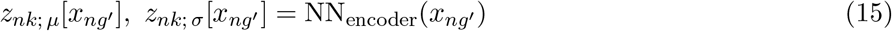

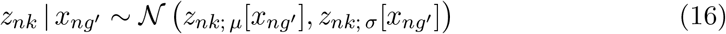

#### Shared projector

To address batch, species and modality differences during horizontal integration, *z*_*nk*_ is concatenated with one-hot encoded batch, species, or modality variables 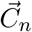 and the shared projector then transforms the concatenated variables into a slightly higher-dimensional space *y*_*np*_. In this study, the shared projector is implemented as a single fully-connected dense layer, without any batch normalization or activation function.

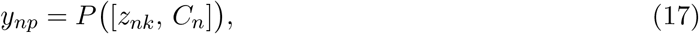

where *p > k*. By explicitly conditioning on covariates, the projector absorbs covariates-related variation, enabling *z*_*nk*_ to capture primarily biologically meaningful structure.

### 5.3 Horizontal integration with multiVIB

In phase 1 of training, the modality-specific translator requires one-to-one or many-to-one correspondences between original and translated features in the form of a mask. For example, a translator for scATAC-seq data that projects features into gene space can be parameterized using a predefined mask that encodes prior regulatory assumptions. Specifically, the mask is constructed under the assumption that chromatin accessibility within a 2 kb window upstream of the transcription start site (TSS) contributes more directly to gene transcription, whereas distal regulatory elements exert comparatively weaker or less direct effects. Accordingly, accessibility peaks falling within this promoter-proximal region are assigned a value of 1 to activate their contribution during the mapping to corresponding gene-level features. Although this masking strategy represents a simplified approximation of cis-regulatory architecture, gene activity matrices derived from such heuristics have been experimentally demonstrated to facilitate effective integration of scATAC-seq and scRNA-seq data [67]. Masked out weights in the translator will be inactive (zero) during phase 1 training. This enables exploration of biologically-informed correlations while preventing the model from learning spurious associations. After initial phase 1 training (100 epochs in this study), we switch to phase 2, lifting the mask constraint from the translator and allowing all weights to be updated. We keep training model for extra epochs (200 epochs in this study), and the second form of the translator should enable the model to capture a broader range of correlations between the original features and the shared space.

For unpaired DNA methylation data to be integrated with horizontal approach, we constructed the a priori mask by assigning a correspondence between each CpG region and its nearest gene, using the criterion that CpG sites falling within 2,000 bp upstream or downstream of a gene body are considered to potentially influence that gene’s expression. This gene-proximal methylation mask reflects the well-established association between promoter-proximal CpG methylation and transcriptional silencing. CpG sites outside this window are assigned a mask value of 0 and their translator weights are not updated during Phase 1 training.

### 5.4 Vertical integration with multiVIB

When cells are jointly profiled to obtain multi-omics data, so that an individual cell has two data modalities measured, we first fit a linear regression model with `sklearn.linear_model.LinearRegression`, which estimates regression coefficients that map features from one modality to the other by minimizing the sum of the squared differences between predicted and observed values. This step discovers correlations between original and translated features, and is used to initialize the linear modality-specific translator. In vertical integration, we primarily skip phase 1 training, acting as though the linear regression results are the output of phase 1, and we directly use the second form as the translator, where the weight matrix is factorized into *A*_*fd*_ and *B*_*dg*_. This reduces the number of trainable parameters. However, users have the option to train using phase 1 training when needed.

### 5.5 Cell embedding through multiVIB

Anchor cells with multimodal profiles are assigned modality-specific embeddings, one per modality. For downstream use, these embeddings are aggregated via simple averaging to obtain a unified representation for each anchor cell. However, depending on the requirements and inductive biases of specific downstream analyses, modality-specific embeddings may provide more appropriate representations than the aggregated embedding. Therefore, we retain both the unified and modality-specific embeddings and allow users to select the representation best suited to their analytical objectives.

In practice, several considerations guide the selection of the final cell embedding. (1) For tasks requiring a unified representation—such as UMAP visualization, clustering, or cross-dataset label transfer—we recommend using the averaged embedding across all available modalities for jointly profiled cells. This aggregation maximizes the contribution of each modality while yielding a single, consistent coordinate per cell. (2) For modality-specific analyses—such as characterizing chromatin accessibility dynamics or gene expression variability—modality-specific embeddings may be preferable, as they preserve signals that could otherwise be attenuated by averaging. (3) For unpaired cells in mosaic integration settings, only modality-specific embeddings are available and are therefore used directly without aggregation. A key implication of foregoing multimodal aggregation is that modality-specific biological variation is more faithfully retained, albeit at the expense of a less harmonized joint representation.

### 5.6 Input data pre-processing

For scRNA-seq data, we took common pre-processing steps by (1) normalizing the total counts to 10,000 scale, (2) log1p-transforming, and (3) standardizing gene features by removing the mean and scaling to unit variance. For scATAC-seq data, we perform a TF-IDF transformation on the accessibility matrix[68]. For a peak *p* in cell *n*, the transformed TF-IDF peak matrix *t*_*np*_ is calculated as:

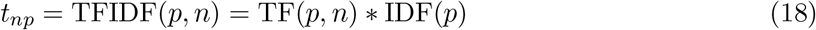

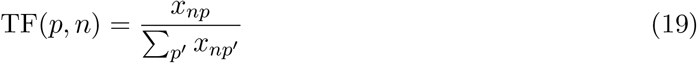

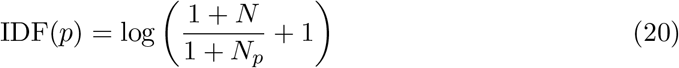

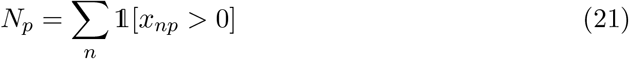

where *x*_*np*_ is the original peak count data, *N* is the total number of cells, and *N*_*p*_ is the number of cells that have peak *p*. We further standardize the transformed accessibility matrix *t*_*np*_ by removing the mean and scaling to unit variance.

For histone modification data, we bin peak features into 10,000bp windows. We log1p-transform the binned data and standardize the histone modification matrix by removing the mean and scaling to unit variance. Since, all input datasets are standardized by removing the mean and scaling to unit variance, we adapt a universal augmentation for all modalities via adding random gaussian noise with mean as 0 and variance as 1 to individual features and cells. We also implement 0.25 feature dropout in the shared encoder to randomly shutdown certain paths from translated features *h*_*ng*_ to latent representation *z*_*nk*_, adding additional augmentations to the input data.

We use scRNA-seq datasets to call out 2000 highly variable genes in each integration task, and these 2000 HVG would serve as the shared gene-centric features. All other modalities will first be translated into the shared 2000 gene feature space through modality specific translator *T* ^(*m*)^.

### 5.7 Evaluation of mouse motor cortex data

For mouse primary motor cortex data, jointly profiled multi-omics data provides multiple data modality measurements per cell. For the purposes of evaluation, we hold out 20% of cells as a test set and use the remaining 80% of cells for training using vertical integration.

### 5.8 Identification of species-specific gene sets

In phase 2 training, the modality translator consists of 2 matrices *A*_*fd*_ and *B*_*dg*_. In cross-species integration (Fig. 4B), we obtain 3 species-specific *A*_*fd*_ matrices and 1 shared 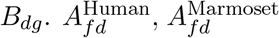, and 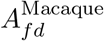 capture species-level regulatory patterns, while *B*_*dg*_ represents a common mapping to the conserved genes across species. We first clustered shared homologous genes into groups using the matrix *B*_*dg*_ and categorized them based on their expression patterns in cell types. Then, we reconstructed a mapping from species-specific genes to shared genes by multiplying *A*_*fd*_ and *B*_*dg*_ to recover *w*_*fg*_ separately for each species. For a given cell type, shared genes that are categorized as being enriched in that cell type were selected. Next, we calculated a z-score of all individual weights of species-specific genes that are linked to shared genes of interest using *w*_*fg*_ in each species, and we defined species-specific genes that have z-scores larger than 2.3263 (equivalent to p-value of 0.01) with at least 5 shared genes as significant in that species. This procedure returns species-specific gene sets that are highly correlated with at least 5 shared homologous genes of interest, which could be further interpreted as having potential regulatory roles in that species, given a cell type of interest.

### 5.9 Evaluation metrics

Luecken *et al*. proposed a comprehensive set of metrics for evaluating single-cell data integration, including cell cycle conservation score and principal component regression (PCR) comparison that operate at the feature level[14]. However, methods benchmarked in this study — including multiVIB — are mostly encoding models, without a generative capacity. Those feature-level metrics are not directly applicable. Computing these feature-level metrics would require an additional decoding step, which could be not standardized across different methods, and we argue that it would introduce confounds into the comparison. Therefore, we restricted our evaluation to embedding-level metrics, which are model-agnostic and directly comparable across all methods.

#### 5.9.1 NMI

We measure the normalized mutual information (NMI) with the sklearn implementation: `sklearn.metrics.normalized_mutual_info_score`. NMI compares the overlap of two clusterings. We used NMI in this study to compare the author-reported cell-type labels with label transfer predictions from another source of dataset. Due to different annotation resolutions of datasets from different studies, NMI scores of 0 or 1 correspond to uncorrelated annotation or a perfect match of annotation resolution, respectively.

#### 5.9.2 ARI

The adjusted rand index (ARI) is implemented with `sklearn.metrics.adjusted_rand_score`. ARI also compares the overlap of two clusterings, and we also included ARI in this study to compare the author-reported cell-type labels with label transfer predictions from another source of dataset. ARI is more sensitive to resolutions of two labels, compared to NMI. If two labels from two different sources have distinct annotation resolutions, the comparison usually returns low ARI score close to 0. Only if resolutions of two labels are very close, the calculation will yield ARI score close to 1.

#### 5.9.3 ASW

We adapted averaged silhouette width score (ASW) from Luecken *et al*. to measure the relationship between the within-cluster distances of a cell and the between-cluster distances of that cell to the closest cluster[14].

To evaluate data integration outputs, we calculated ASW scores with (1) cell type labels (“ASW cell type”) and (2) combined cell type labels and modality labels to measure modality mixing (“ASW modality”).

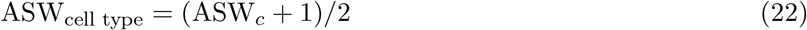

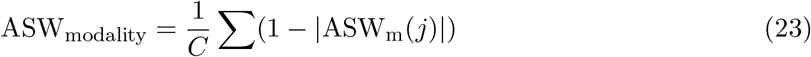

where ASW_*c*_ is a classical definition of silhouette width score computed using `sklearn.metrics.silhouette_score`. *c* denotes the set of cell type labels. For each cell type *j*, ASW_m_(*j*) is calculation of classical silhouette width score computed using modality label. Then, the final ASW_modality_ is the average score of 1 − |ASW_m_(*j*)| across all cell types *C*.

#### 5.9.4 Batch and modality mixing score

We adapted an entropy-based calculation to measure mixing of batch or modality from Xiong *et al*.[69]. Batch or modality mixing score measures a regional mixing of cells from different batches or modalities, with a high score suggesting cells from different sources are well mixed. This score is computed as

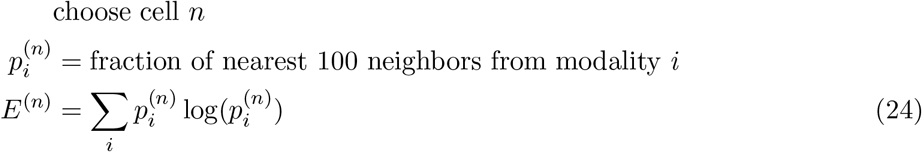

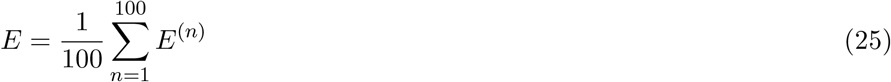

In our calculation, a given region is represented by a random sample of 100 cells from each batch or modality. 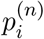 is the fraction of cells from batch or modality *i* in the 100 nearest neighbors of cell *n*. This process is repeated 100 times to get an averaged batch or modality mixing score *E*.

#### 5.9.5 Graph connectivity

Graph connectivity [14] assesses whether cells belonging to the same biological cell type remain mutually reachable within the *k*-nearest neighbor (kNN) graph constructed on the integrated embedding. For each cell type *c*, the metric identifies the subgraph induced by all cells of that type and computes the fraction of those cells contained in the largest connected component. Formally, the graph connectivity score for cell type *c* is defined as

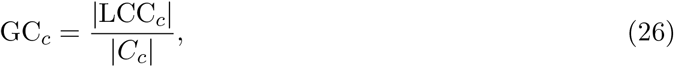

where |LCC_*c*_| is the size of the largest connected component and |*C*_*c*_| is the total number of cells of type *c*. The overall graph connectivity score is the mean of GC_*c*_ across all cell types, weighted by cell type frequency. A score of 1 indicates that all cells of every cell type form a single fully connected subgraph in the integrated space, reflecting complete biological coherence; a score approaching 0 indicates severe fragmentation, where cells of the same type are scattered into disconnected islands in the embedding. Unlike cluster-based metrics such as ARI or NMI, graph connectivity operates directly on the neighborhood graph topology and is therefore sensitive to local embedding structure rather than global cluster assignments, making it particularly informative for detecting cases where integration preserves broad cluster identity but fails to connect transitional or rare cell states within each type.

#### 5.9.6 Label transfer

Label transfer evaluates cross-modal mapping by assessing how accurately cell-type annotations from one modality can be predicted in another. We quantified label transfer performance using overall accuracy and macro F1 score. These two metrics are used when the source data and target data has the same annotation resolution. Accuracy presents an overall prediction accuracy, regardless of cell types. On the other hand, macro F1 shows an average prediction across cell types. Thus, a low macro F1 score but high overall accuracy would indicate inaccurate prediction for minor cell types.

#### 5.9.7 Local annotation enrichment score

To evaluate modality mixing at a local neighborhood scale, we designed the metric, local annotation enrichment score (LAES)[13]. For each cell type, we calculate the *k*-nearest neighbors of individual cells in the unified latent space *z*_*nk*_. The per-cell LAES_*n*_ and mean LAES for the cell-type population are calculated as:

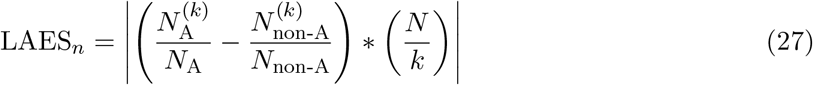

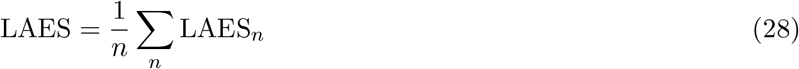

where *n* is the index of a cell in modality 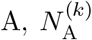 is the number of cells in modality A among the *k*-nearest neighbors of cell 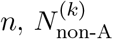, is the number of cells in all other modalities among the *k*-nearest neighbors of cell *n, k* is the number of nearest neighbors used in the computation, and *N* is the total number of cells of a certain cell type combining all modalities.

LAES estimates the local discrepancy between density of points from two or more sources: in this case, cells from different modalities. LAES close to 0 indicates that samples from these sources may come from the same distribution, which suggests good modality mixing in the latent space *z*_*nk*_.

#### 5.9.8 Average score for method ranking

To report a method ranking in benchmarking, we first ranked methods in individual evaluation metrics. Over 4 evaluation categories, we averaged the ranking index of each method in each evaluation category. We then calculated the final ranking index *R* of method *i* as below:

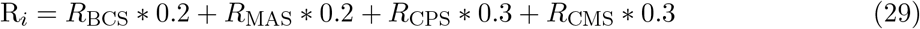

where BCS stands for batch correction score, MAS for modality alignment score, CPS for cell-type preservation score, and CMS for cross-modal mapping score.

Our averaged ranking gives higher weights on biological information-related metrics (preservation of biological variation and cross-modal mapping).

## Data Availability

10X Multiome PBMC dataset was accessed via the NeurIPS 2021 Datasets and Benchmarks track. Other human PBMC and bone marrow datasets used in this study could be found through the original studies, with Gene Expression Omnibus (GEO) (RRID:SCR 005012) accession numbers GSE139369, GSE129785, and GSE216005, respectively. Processed batch correction benchmark datasets, including single-cell RNA-seq immune cell data and single-cell ATAC-seq mouse brain cell atlas data, were obtained from benchmarking study by Luecken *et al*.[14], with link at https://doi.org/10.6084/m9.figshare.12420968. Human lung cell atlas data can be downloaded via cellxgene (RRID:SCR 021059) (https://cellxgene.cziscience.com/collections/6f6d381a-7701-4781-935c-db10d30de293). Cross-species primary motor cortex data of human and mouse were generated by two previous BRAIN Initiative studies[46, 55]. Processed version of both datasets can be downloaded via cellxgene (https://cellxgene.cziscience.com/collections/ae1420fe-6630-46ed-8b3d-cc6056a66467 and https://cellxgene.cziscience.com/collections/d17249d2-0e6e-4500-abb8-e6c93fa1ac6f) Preprocessed cross-species basal ganglia datasets can also be found at https://alleninstitute.github.io/HMBA_BasalGanglia_Consensus_Taxonomy/. Previously collected mouse primary motor cortex datasets of RNA-seq, DNA methylation, and ATAC-seq under BICCN initiative can be found under NeMO https://nemoarchive.org/ RRID:SCR_016152. Preprocessed versions of these datasets are also available at https://cellxgene.cziscience.com/datasets. Joint profiling mouse primary motor cortex data of gene expression and chromatin accessibility through ISSAAC-seq was deposited at ArrayExpress (RRID:SCR 002964) under the accession number E-MTAB-11264. Joint profiling mouse primary motor cortex data with histone modifications through Pair-tag was deposited at GEO (RRID:SCR 005012) under accession number GSE152020.

## Code Availability

multiVIB (RRID:SCR_027589) code is available at https://github.com/broadinstitute/multiVIB. Tutorials using multiVIB for data integration are also provided in the multiVIB GitHub repository. The public release of multiVIB code is also available at Zenodo (https://doi.org/10.5281/zenodo.19926776)[70].

## Acknowledgments

This publication was supported by and coordinated through the Brain Initiative Cell Atlas Network (BICAN). The study was supported by NIH grant RF1MH133777 and the Broad Institute Data Sciences Platform.

## S Supplemental Figures

**Supplementary Figure 1:**
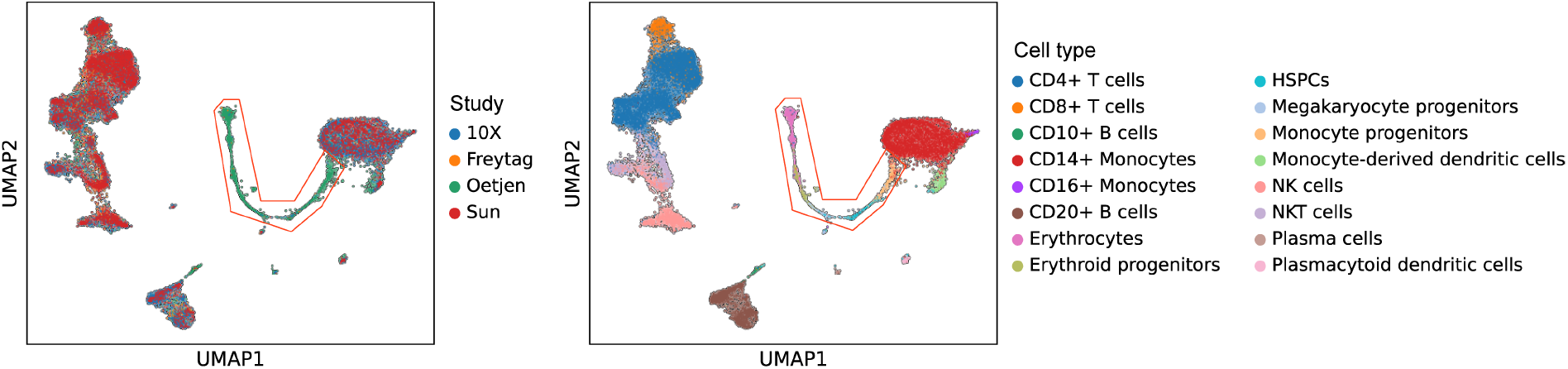
Learned representations preserve cell-type structure without over-correcting batch effects in PBMC and BMMC data. UMAP visualization of integrated cell embeddings colored by study of origin (left) and annotated cell type (right). Four studies (10X, Freytag, Oetjen, and Sun) are jointly embedded. The left panel demonstrates effective batch mixing within each cell-type cluster, while the right panel confirms that biologically distinct populations remain well-separated in the shared latent space. The red box highlights a region of particular interest where multiple cell types converge, illustrating that integration does not collapse transcriptionally distinct populations into cell types of the other 3 studies.

**Supplementary Figure 2:**
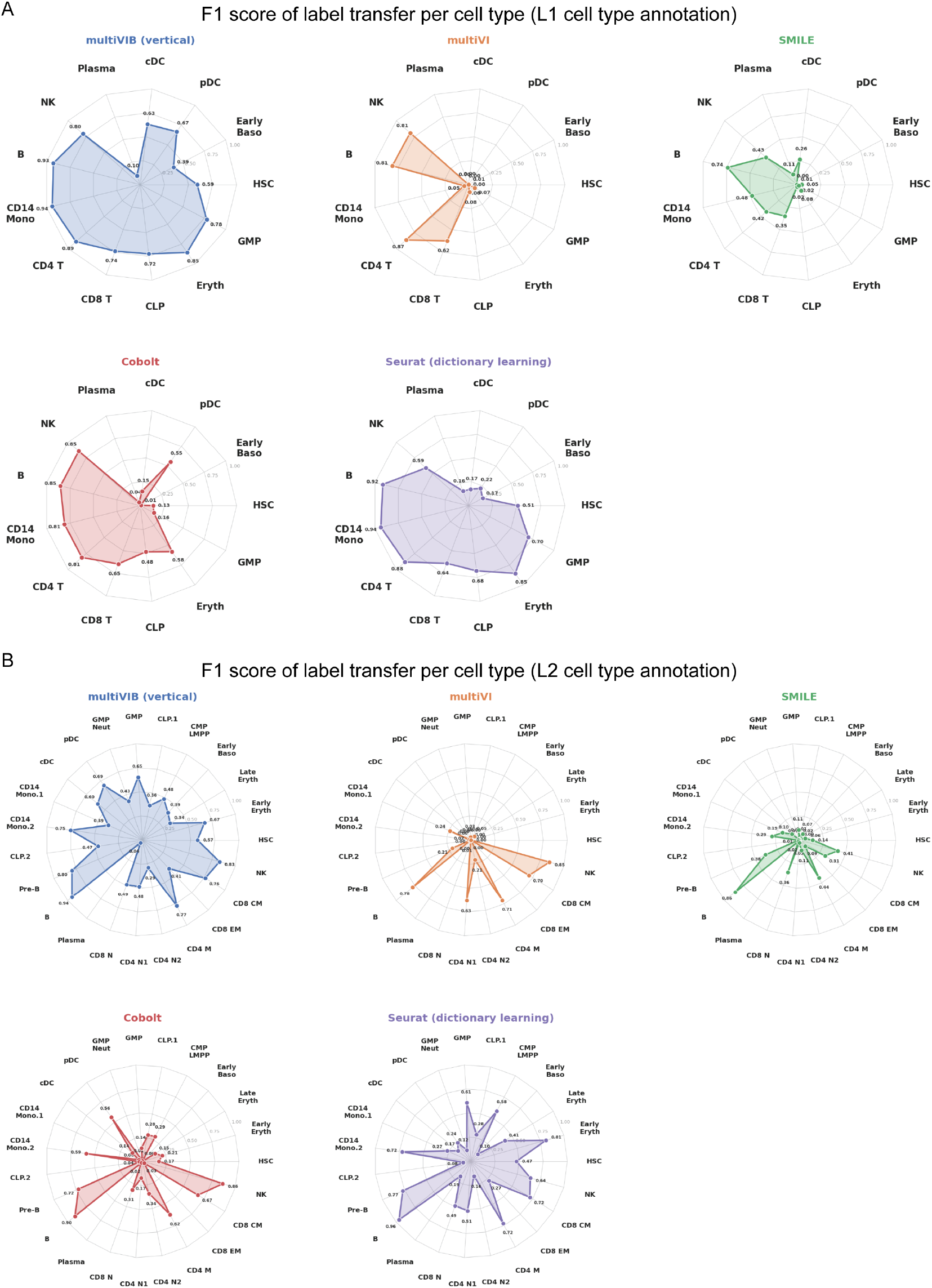
Per-cell-type F1 scores for cross-modal label transfer in vertical integration. Radar charts displaying per-cell-type F1 scores for cross-modal label transfer, comparing multiVIB (vertical) against multiVI, SMILE, Cobalt, and Seurat (dictionary learning) at (A) coarse (L1) and (B) fine-grained (L2) cell type annotation resolutions. Each axis represents a distinct cell type, and radial distance indicates F1 score (range: 0–1.0). multiVIB (vertical) achieves consistently high and balanced F1 scores across the majority of cell types at both annotation levels, whereas competing methods exhibit pronounced imbalances — performing well on abundant cell types such as CD14 monocytes and B cells while failing on rare populations including HSC, GMP, pDC, and early progenitor subtypes. The advantage of multiVIB is particularly evident at L2 resolution, where the increased number of fine-grained cell type categories amplifies performance gaps between methods.

**Supplementary Figure 3:**
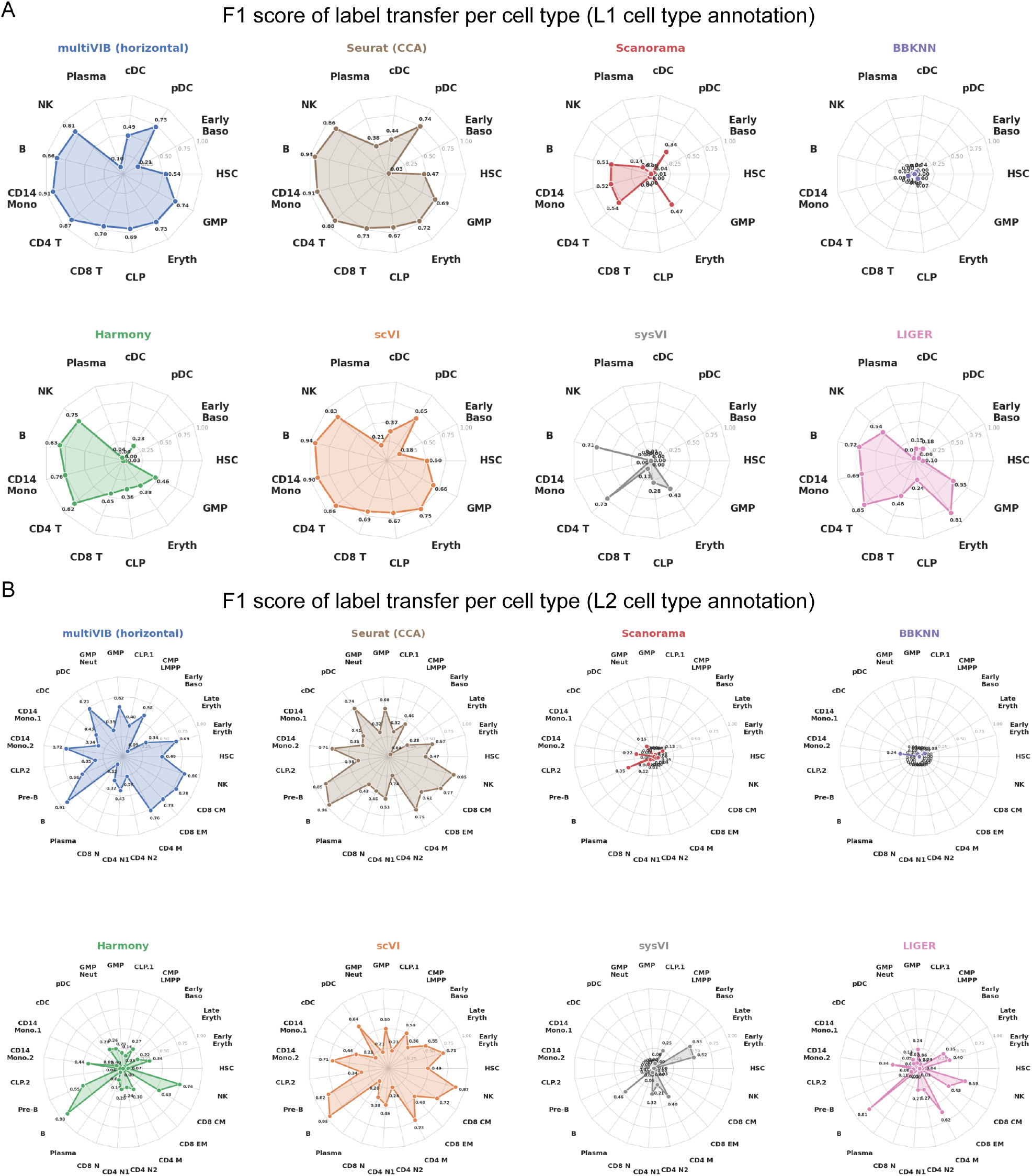
Per-cell-type F1 scores for cross-modal label transfer in horizontal integration. Radar charts displaying per-cell-type F1 scores for cross-modal label transfer, comparing multiVIB (horizontal) against Seurat (CCA), Scanorama, BBKNN, Harmony, scVI, sysVI, and LIGER at (A) coarse (L1) and (B) fine-grained (L2) cell type annotation resolutions. Each axis represents a distinct cell type, and radial distance indicates F1 score (range: 0–1.0). multiVIB (horizontal) demonstrates the most balanced performance profile across cell types at both annotation levels. Methods such as BBKNN and Scanorama exhibit severely collapsed radar profiles, reflecting near-zero F1 scores for the majority of cell types. At L2 resolution, multiVIB, scVI, and Seurat (CCA) maintain broader and more uniform coverage across the expanded set of fine-grained cell type categories compared to all other evaluated methods, underscoring the advantage of multiVIB for cross-modality label transfer in horizontal integration settings.

**Supplementary Figure 4:**
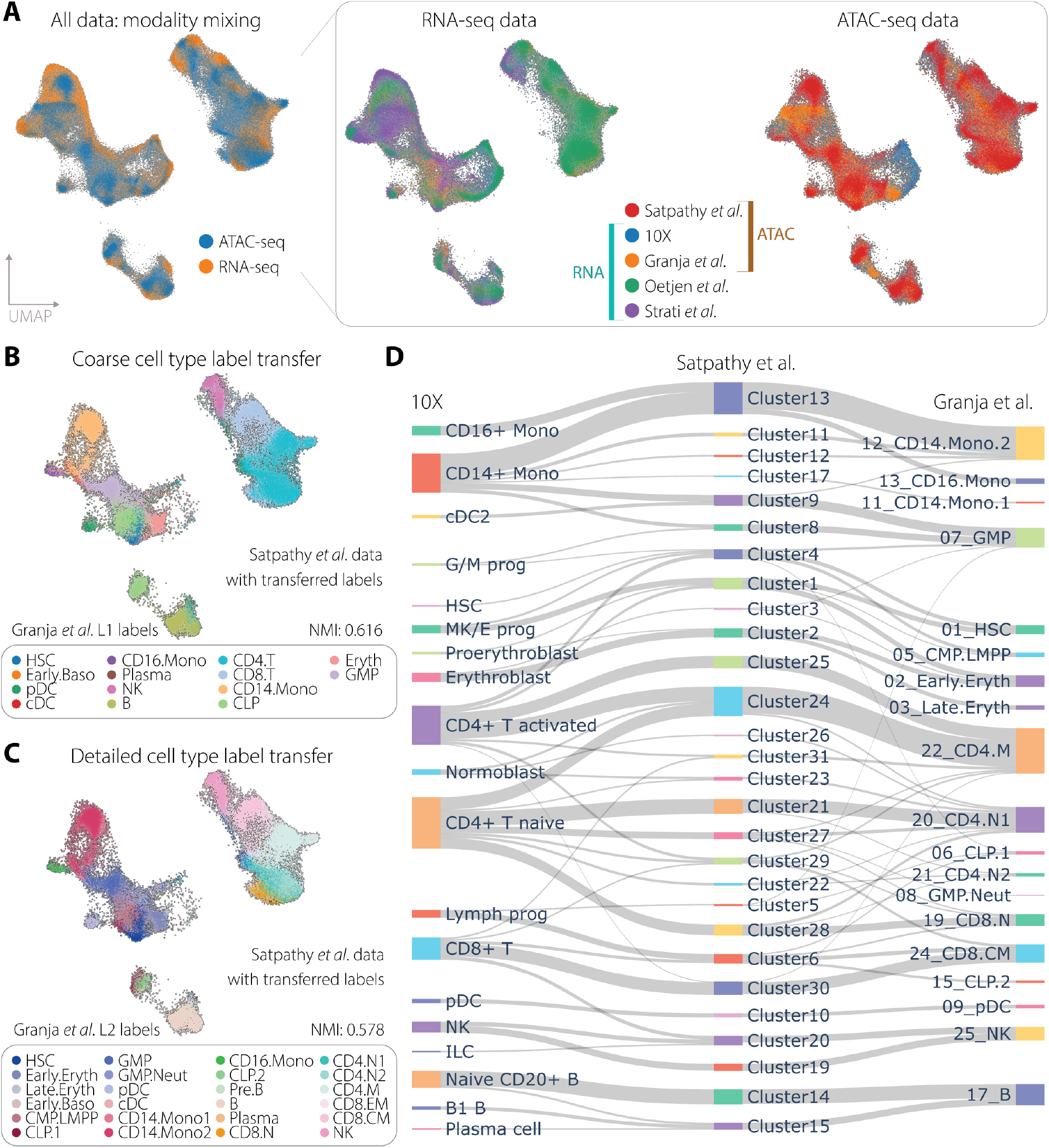
multiVIB enables label transfer and cell querying. (A) UMAP visualization of multi-study human PBMC data integration via multiVIB (vertical). Cells are colored according to modality source (left), batch ID of gene expression modality (middle), and batch ID of chromatin accessibility modality (right). (B, C) Label transfer from Granja*et al*. scRNA-seq data to Satpathy*et al*. scATAC-seq data. Transferred labels are evaluated with author-reported clustering label using NMI. (B) Coarse cell type labels (L1) were transferred. (C) Detailed, granular cell type labels (L2) were transferred. (D) Mapping cells of Satpathy*et al*. scATAC-seq data to 10X and Granja*et al*. scRNA-seq data. The top 15 similar cells in 10X and Granja*et al*. scRNA-seq data for each cell in Satpathy*et al*. scATAC-seq data were retrieved using the multiVIB latent space. Majority voting is used to identify the most similar cell type for each cell of Satpathy*et al*. scATAC-seq data. The gray connectors link cells from the query Satpathy*et al*. scATAC-seq data to the other two reference datasets.

**Supplementary Figure 5:**
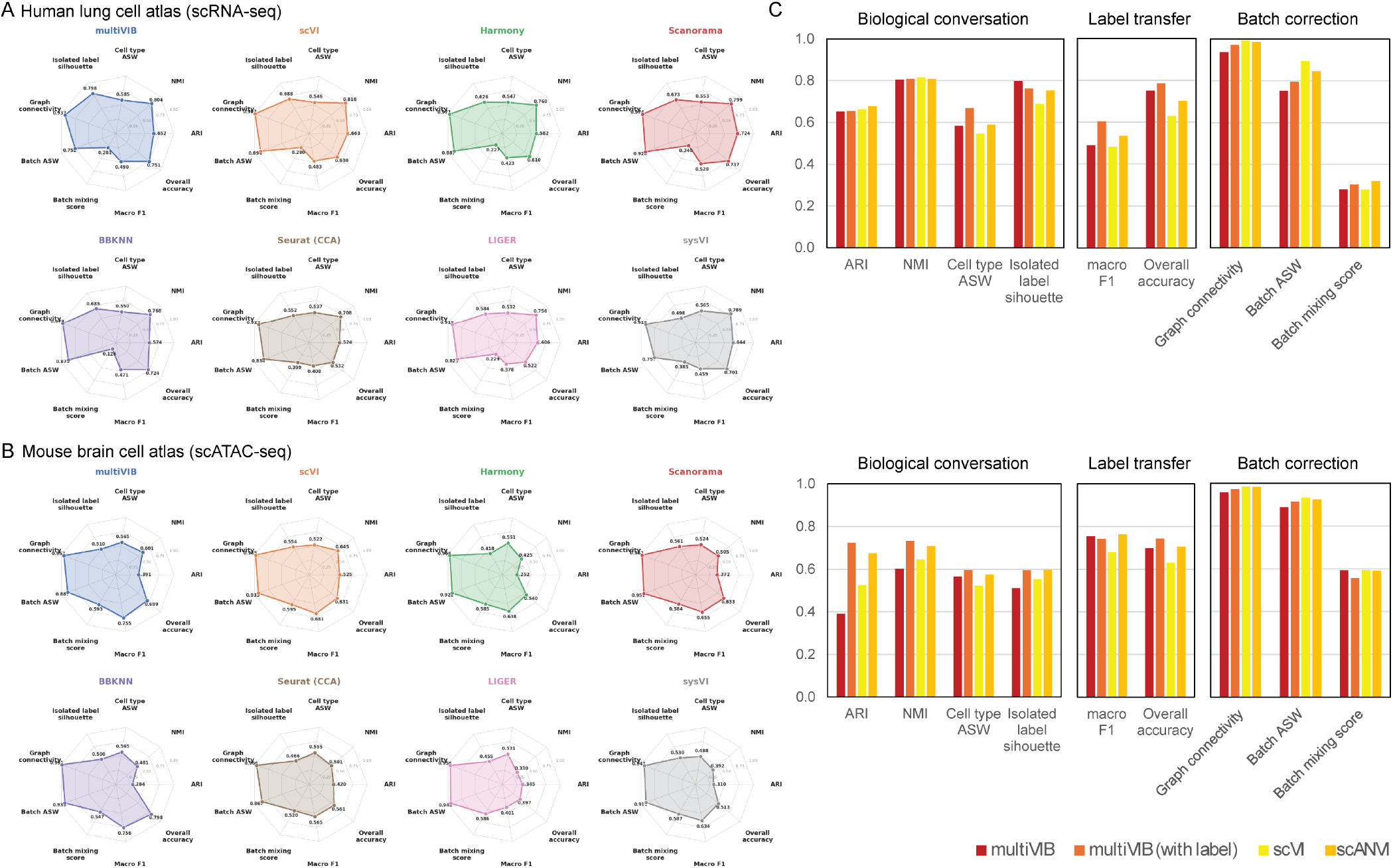
Comprehensive benchmarking of multiVIB against state-of-the-art integration methods across scRNA-seq and scATAC-seq datasets. (A–B) Radar charts comparing multiVIB with seven integration methods — scVI, Harmony, Scanorama, BBKNN, Seurat (CCA), LIGER, and sysVI — across nine metrics spanning biological conservation (ARI, NMI, cell type ASW, isolated label silhouette), batch correction (graph connectivity, batch ASW, batch mixing score), and label transfer (macro F1, overall accuracy), evaluated on (A) the human lung cell atlas (scRNA-seq) and (B) the mouse brain cell atlas (scATAC-seq). Metrics follow the benchmarking framework established by Luecken *et al*.[14]. Larger radar area indicates stronger overall performance. (C) Bar charts directly comparing unsupervised methods (multiVIB and scVI) against semi-supervision methods (multiVIB with label and scANVI) across the three metric categories — biological conservation, label transfer, and batch correction — for the human lung cell atlas (top) and mouse brain cell atlas (bottom). multiVIB trained with partial cell type labels achieves performance comparable to or exceeding scANVI in both datasets. All metrics are computed in the latent space. Both biological conservation scores (ARI, NMI, Cell type ASW, Isolated label silhouette, Macro F1, and Overall accuracy) and data alignment scores (Graph connectivity, Batch mixing score, and Batch ASW) are all annotated as higher is better.

**Supplementary Figure 6:**
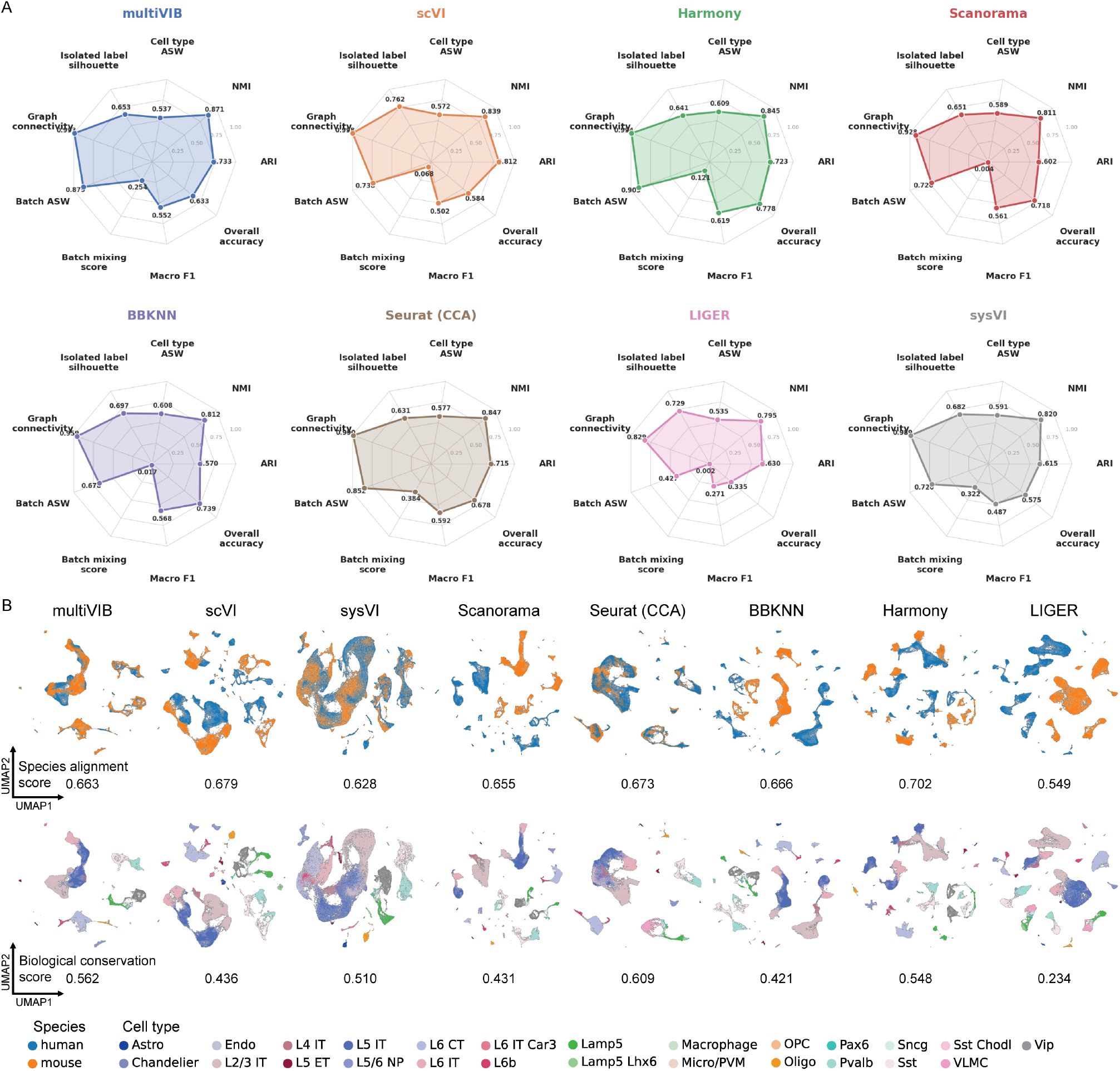
Benchmarking of horizontal integration methods on primary motor cortex data from human and mouse. (A) Radar charts comparing multiVIB against seven horizontal integration methods — scVI, Harmony, Scanorama, BBKNN, Seurat (CCA), LIGER, and sysVI — across nine metrics spanning biological conservation (ARI, NMI, cell type ASW, isolated label silhouette), species alignment (graph connectivity, batch ASW, batch mixing score), and label transfer (macro F1, overall accuracy). Larger radar area indicates stronger overall performance. (B) UMAP projections of the integrated latent representations for all eight methods, colored by species (top row; human: blue, mouse: orange) and annotated cell type (bottom row). Species alignment scores and biological conservation scores are reported below each UMAP pair, respectively. multiVIB, aside with sysVI and Seurat (CCA), achieves the highest species alignment scores while maintaining competitive biological conservation, demonstrating the capacity of multiVIB to simultaneously align transcriptomic representations across species and preserve fine-grained cell type structure in primary motor cortex data. Both biological conservation scores (ARI, NMI, Cell type ASW, Isolated label silhouette, Macro F1, and Overall accuracy) and data alignment scores (Graph connectivity, Batch mixing score, and Batch ASW) are all annotated as higher is better.

**Supplementary Figure 7:**
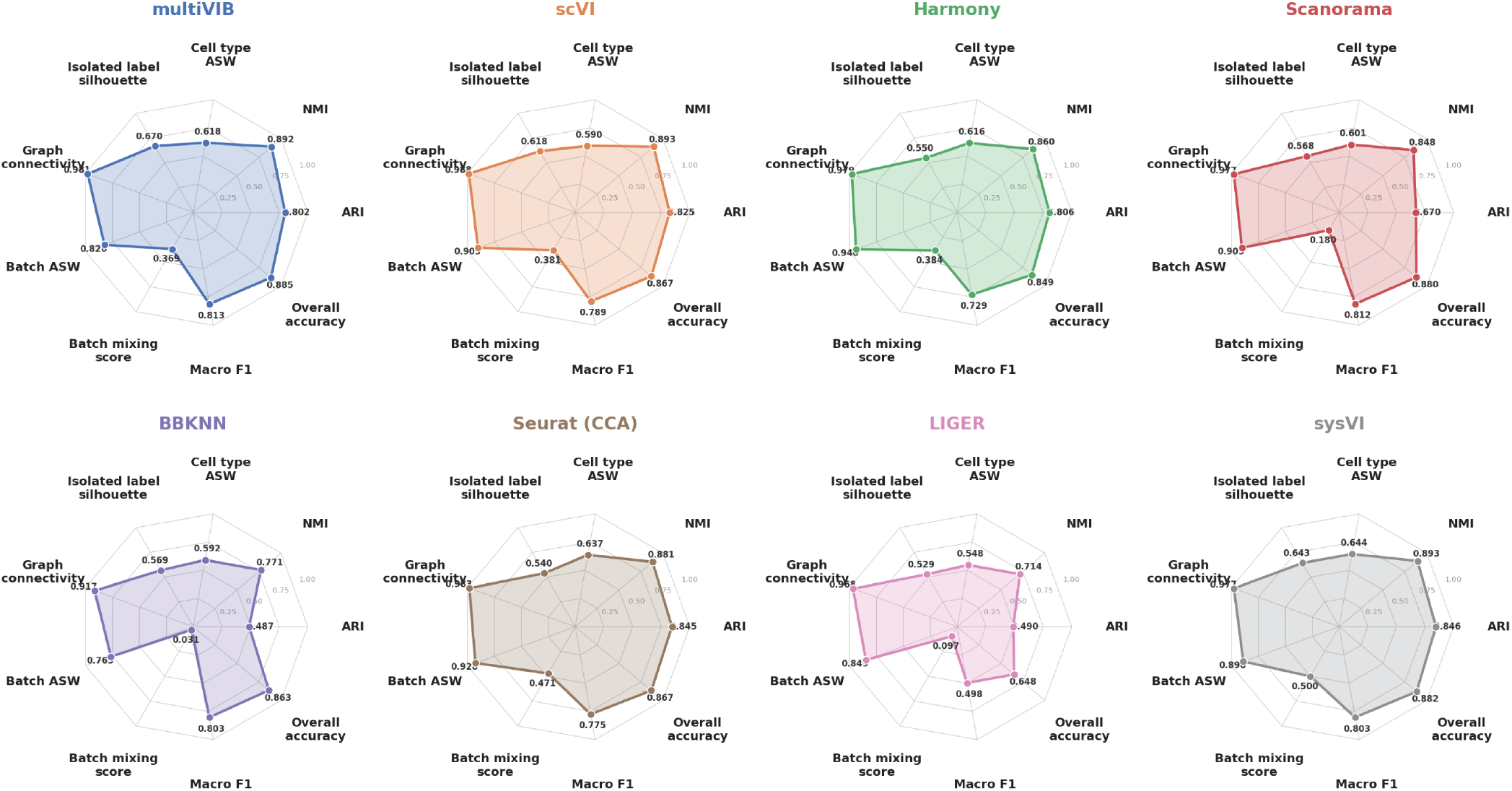
Quantitative benchmarking of horizontal integration methods on cross-species basal ganglia dataset. Radar charts comparing multiVIB against seven horizontal integration methods — scVI, Harmony, Scanorama, BBKNN, Seurat (CCA), LIGER, and sysVI — across nine metrics spanning biological conservation (ARI, NMI, cell type ASW, isolated label silhouette), species alignment (graph connectivity, batch ASW, batch mixing score), and label transfer (macro F1, overall accuracy). Harmonized cross-species cell type annotations from Johansen et al.[3] were used as ground truth for all clustering and label transfer metrics. Larger radar area indicates stronger overall performance. multiVIB achieves a balanced and competitive profile across both biological conservation and species alignment metric categories, establishing it as one of the leading methods for cross-species transcriptomic integration of basal ganglia data. All metrics (ARI, NMI, Cell type ASW, Isolated label silhouette, Macro F1, and Overall accuracy, Graph connectivity, Batch mixing score, and Batch ASW) are all annotated as higher is better.

**Supplementary Figure 8:**
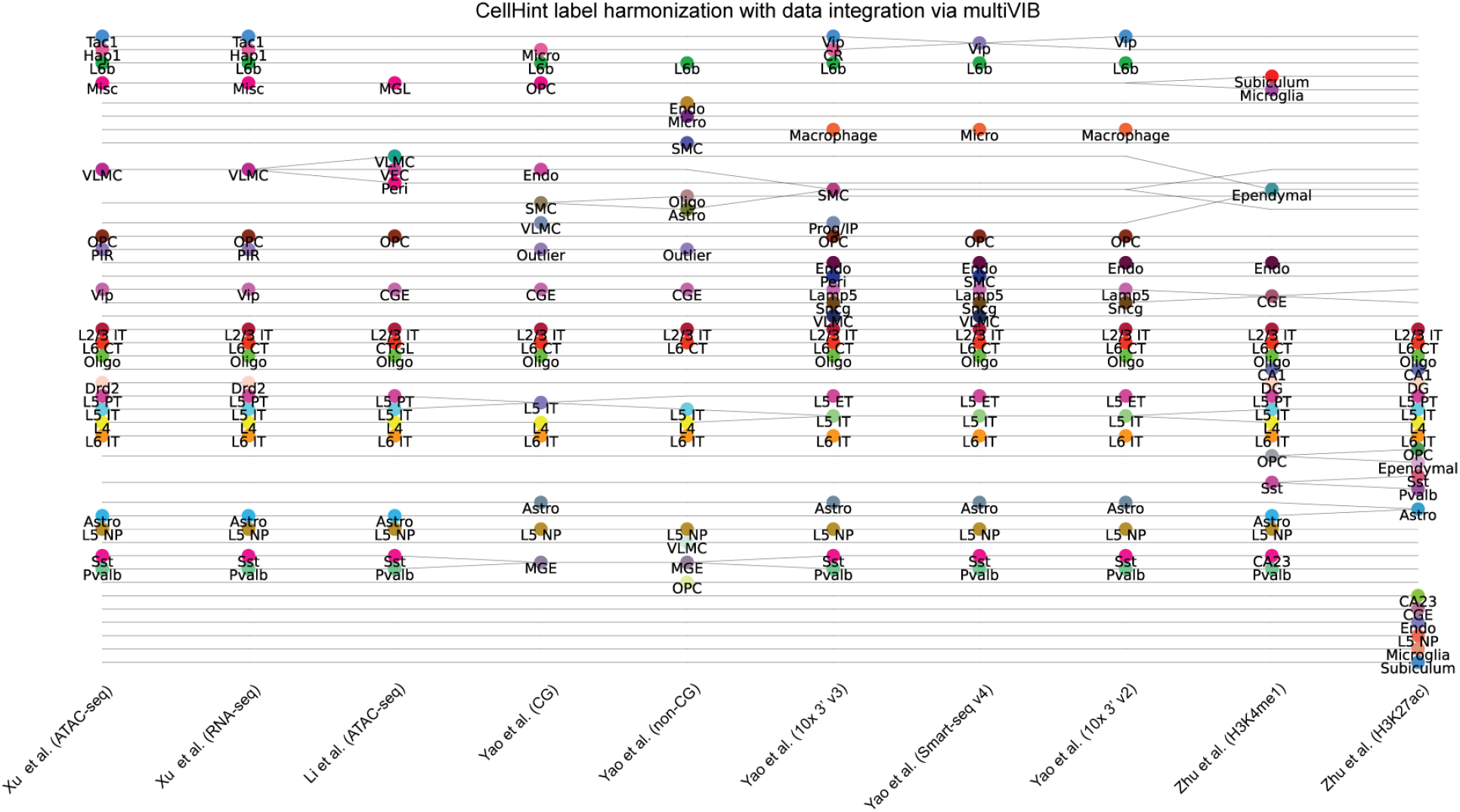
CellHint annotation harmonization of mouse primary cortex data with multiVIB cell embedding. Mapping cell types across modalities is computed using the tool CellHint. CellHint takes cell embeddings and cell type labels to harmonize cell type labels across datasets. Here, we applied CellHint to map the consensus cell type across modalities. For cell embeddings, we used the unified cell embeddings learned with multiVIB. Using multiVIB cell embedding CellHint accurately mapped cell types that have the same identity across modalities, with exception that some cell types of H3K27ac data modality cannot be accurately matched with other modalities.

**Supplementary Figure 9:**
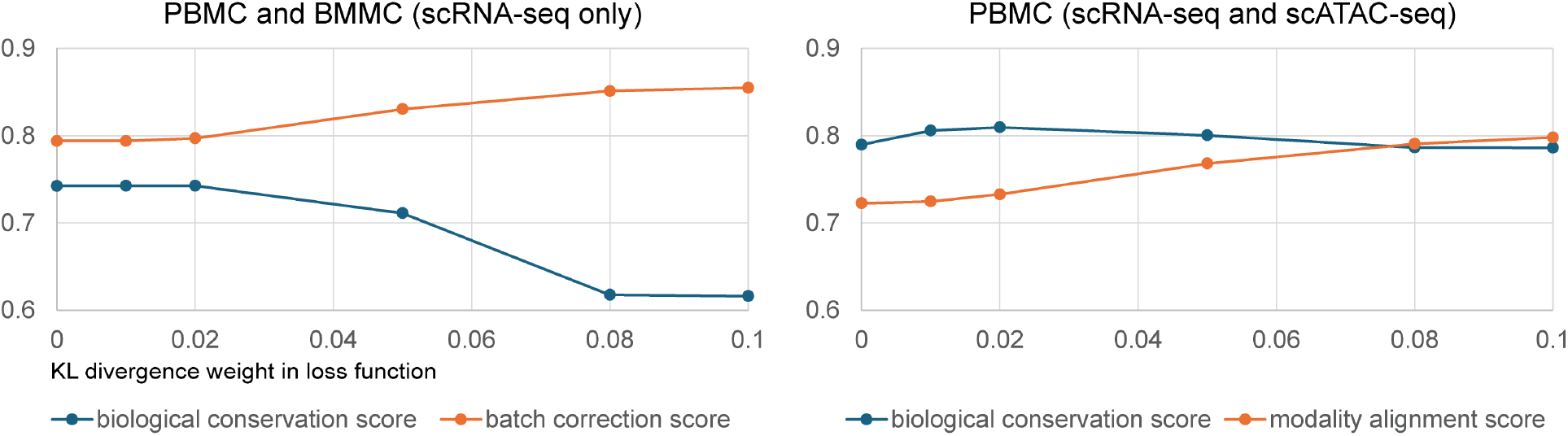
Effect of KL divergence weight on the trade-off between biological conservation and dataset alignment. Line plots showing how varying the KL divergence weight in the multiVIB loss function affects biological conservation score and dataset alignment score, evaluated on two example datasets: PBMC and BMMC (scRNA-seq only, horizontal integration; left) and PBMC (paired scRNA-seq and scATAC-seq, vertical integration; right). In the horizontal integration setting, increasing the KL divergence weight monotonically improves batch correction score but leads to a pronounced decline in biological conservation score beyond a weight of 0.02, indicating over-regularization that collapses biologically distinct cell type structure. In the vertical integration setting, both biological conservation and modality alignment scores remain relatively stable at low weights before converging at higher values, reflecting a more gradual trade-off. Across both settings, a KL divergence weight in the range of 0.02–0.05 achieves a favorable balance between preserving biological variation and promoting cross-dataset or cross-modality alignment.

